# An *in situ* cut-and-paste genome editing platform mediated by CRISPR/Cas9 or Cas12a

**DOI:** 10.1101/2022.03.30.486486

**Authors:** Ping Jiang, Kevin M. Kemper, Kai-Ti Chang, Cheng Qian, Yulong Li, Liying Guan, Peter van Hasselt, Salvatore J. Caradonna, Randy Strich

**Affiliations:** Department of Molecular Biology, Rowan University School of Osteopathic Medicine, Stratford, New Jersey, USA; Euprotein Inc. North Brunswick Township, New Jersey, USA; College of Life Science, Peking University, Beijing, China; Department of Metabolic and Endocrine Disease, University Medical Center Utrecht, Utrecht, Netherlands

**Keywords:** *in situ* cut-and-paste, genome editing, CRISPR, Cas9, Cas12a, mouse genome, human genome, double strand break, DNA repair

## Abstract

Recombinant DNA technology mediated by restriction enzymes and ligases allows *in vitro* manipulation of a DNA segment isolated from the genome. Short overhangs generated by restriction enzymes facilitate efficient pasting together a DNA sequence and a vector. We adopted this recombinant DNA strategy to develop an *in vivo* recombinant-genome genome editing approach. Using the programmable endonuclease Cas9 or Cas12a as a restriction enzyme, we devised an *in situ* <u>c</u>ut-<u>a</u>nd-<u>p</u>aste (iCAP) genome editing method that was tested in both mouse germline and human cell line platforms. Mouse gene loci *Slc35f2* and *Slc35f6* were each edited with in-frame insertion of a large APEX2-Cre cassette and concurrent FRT3 insertion at a second location providing proof of principle for the iCAP method. Further, a *de nova* single nucleotide mutation associated with *MED13L* syndrome was efficiently corrected in patient cells. Altogether, the iCAP method provides a single genome editing platform with flexibility and multiutility enabling versatile and precise sequence alterations, such as insertion, substitution, and deletion, at single or multiple locations within a genomic segment in mammalian genomes.

## INTRODUCTION

Powered by restriction endonucleases, recombinant DNA technologies allow a segment of DNA to be isolated from genomes and then recombined with a vector *in vitro* (Meselson and Yuan, 1968; Cohen et al.,1973). In contrast, it has been more difficult to recombine a segment of *in vitro* restructured DNA back to its original location in a genome, particularly in a mammalian genome. For decades, the only effective approach to introduce *in vitro* reconstructed DNA back into its native location in a mammalian genome is gene-targeting by homologous recombination in mouse embryonic stem cells (mESC) (Thomas and Capecchi, 1987). Generating gene-targeted mESC and animals is a lengthy process involving specialized technical skills and often requires core facility services.

Discovery and application of the CRISPR/Cas9 system (Jinek et al., 2012; Cong et al., 2013) revolutionized targeted DNA editing in mammalian genomes as the system is applicable to virtually all cell types including zygotes. Although highly efficient in targeted gene disruption (knockout) due to aberrant double strand break (DSB) repair, it is still a challenge to precisely alter the DNA sequence at targeted gene loci using orthodox CRISPR/Cas system (Doudna, 2020). New methodologies reported to improve targeted genome editing using CRISPR/Cas9 (Yamamoto and Gerbi, 2018; Pawelczak et al., 2018) generally fall into two categories. The first kind requires DSB induction by CRISPR/Cas9 to initiate homology-directed DNA repair (HDR) when a DNA replacement template (dRT) is available (Cong et al., 2013; Wang et al., 2013). The other class uses engineered Cas variants, such as Cas9 (D10A) nickase, dCas base editors, Cas9 (D10A) nickase-reverse transcriptase, or CRISPR-associated transposases (Ran et al., 2013; Mali et al., 2013; Kim et al., 2017; Gaudelli et al., 2017; Anzalone et al., 2019; Strecker et al., 2019; Klompe et al., 2019), and does not induce DSB. Each of the improved methods has limitations such as a single utility as base editors, limited sequence size for targeted insertion using DNA donors composed of short or long single-stranded oligodeoxynucleotides (ssODNs or lssODNs) (Cong et al., 2013; Li et al., 2017; Miura et al., 2018; Wang et al., 2013; Yang et al., 2013; Yoshimi et al., 2016), or requirements of unique endogenous genomic sequence structures (Iyer et al., 2019; Yokouchi et al., 2020).

As more single-utility improved methods are added to the genome editing toolbox, we investigated whether we could develop a single approach with multiutility and flexibility to achieve different genome editing schemes with precision by simply using native or enhanced CRISPR/Cas effectors. Inspired by the simplicity and flexibility of “cutting” and “pasting” DNA sequences in molecular cloning, we devised an *in situ* <u>c</u>ut-<u>a</u>nd-<u>p</u>aste (iCAP) genome editing platform mediated by Cas9 or Cas12a (Cpf1). Using this approach, we successfully generated genome edited mice that carry a large in-frame inserted expression cassette in *Slc35* gene loci or corrected a single nucleotide mutation in *MED13L* syndrome patient cells, demonstrating its feasibility as a single genome editing platform enabling a wide array of genome engineering schemes.

## RESULTS

### Design of iCAP using Cas9 for mouse genome editing

The mode of iCAP action (Fig. 1a) involves in-nucleus (*in situ*) simultaneous Cas effector (as a “restriction enzyme”) cleavage at two gRNA target sites on the genome as well as at two gRNA target sites on the dRT. Cutting on the genome (as a “vector”) excises an intervening sequence, a genomic segment referred to as the genomic target segment (GTS). Cleavage at two gRNA target sites positioned near the two ends of a dRT fragment releases an “insert” that is an *in vitro* restructured GTS containing precisely edited sequences and thereby referred to as the edited isogenic fragment (EIF). Following a successful “cutting” operation, two distant genomic DSB ends and two EIF’s DNA ends are exposed. The end sequences at the corresponding genomic and EIF ends are defined by design for facilitating end joining mediated by DNA repair machinery with no or minimal end modifications (Lieber, 2010; Chang et al., 2017). Joining of the “insert” with two distant genomic DSB ends results in “paste” of the EIF into the gapped genome.

**Figure 1.**
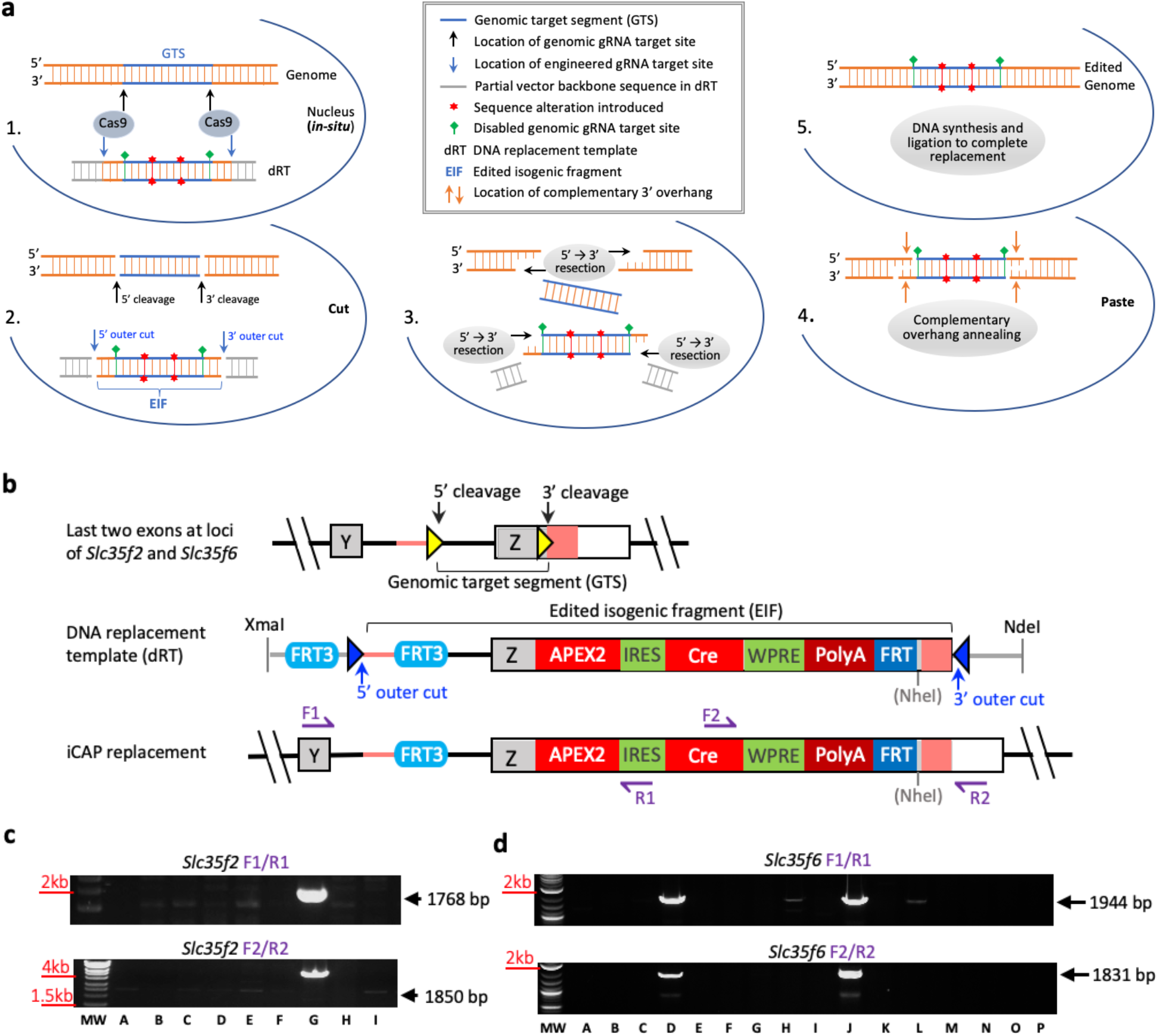
Overview of iCAP genome editing and experimental testing in mouse genome editing. **(a)** Depicted strategy of iCAP genome editing mediated by CRISPR/Cas9. (**b**) Precisely concurrent insertion of exogenous sequences at two locations in each of two mouse gene loci by iCAP. Schematic of the gene region spanning last two exons in *Slc35f2* and *Slc35f6* gene loci (top). Two gRNA target sequences (yellow arrowhead) where two insertions are intended are Cas9 cleavage sites for excision of GTS. Salmon line and box, overhang substrate sequence (OSS). Schematic of Xmal-NdeI restricted dRT construct (middle). Cas9 cleavages at engineered gRNA target sites (blue arrowhead) release EIF that is GTS with precise insertion of the FRT3 site and expression cassette and flanked by 40-bp OSS (salmon line and box). Grey line, vector backbone sequence; grey NheI, restriction site lost. Predicted iCAP edited gene locus (bottom). Cas9 cleavages at the gene locus and dRT in nucleus (*in-situ*) excise both GTS and EIF exposing defined DNA ends, OSS. In end-resection mediated DNA repairs, 5’→3’ end resection within OSS creates complementary 3’ overhangs at corresponding genomic and EIF ends for annealing and ligation resulting in iCAP replacement of GTS with EIF. F1/R1 and F2/R2, locations of primer pairs for genotyping. (**c)** Genotyping results of 9 animals (A-J) indicates Animal G as a candidate carrying iCAP edited *Slc35f2* gene locus. (**d**) Genotyping results of 16 animals (A-P) indicates Animals D and J as candidates carrying iCAP edited *Slc35f6* gene locus.

To test the feasibility of the iCAP approach, we produced reporter-Cre mice that carry one of three edited alleles, *Slc35f2, Slc35f6,* or *P2y14* (*P2ry14*). The modified alleles contain (i) a 48-bp FRT3 site inserted in intron 7-8 of *Slc35f2*, in intron 5-6 of *Slc35f6* and in intron 1-2 of *P2y14*, and (ii) a 3.7 kbp APEX2-IRES-CRE-WPRE expression cassette placed in-frame immediately 3’ of the codon for the last amino acid in the C-terminus in exon 8 of *Slc35f2* and in exon 6 of *Slc35f6* (Fig. 1b and S1a-b), respectively, or the similar sized EGFP-IRES-CRE-WPRE expression cassette inserted in-frame immediately 3’ of the codon for the last amino acid in exon 2 of *P2y14* (Fig. S1c and S4a). The in-frame insertion of the expression cassette, that contains stop codons and hGHpolyA, would eliminate function utilities of the endogenous 3’ non-coding sequences in the last exons. Such precise gene editing for a mammalian genome remains to be a technical challenge to accomplish using currently available genome editing methods.

To employ iCAP strategy to edit the alleles as described above, we first identified upstream (5’) gRNA target sequences in these three introns as the 5’ cleavage site and downstream (3’) gRNA target sequences near the stop codons as the 3’ cleavage site (Fig. 1b, S1a-c, and S4a). These gRNA targets were validated for CRISPR/Cas9 induced DSBs to excise the intervening sequences, GTS, in an *in-vitro* cleavage assay. We next used the GTS sequences bearing the genomic gRNA target sites as a portion of isogenic DNA to build EIF-containing dRTs. To prevent cleavage at the genomic gRNA target sites present on the dRTs, we disrupted these sites by inserting a FRT3 site into the 5’ gRNA target sequence and the expression cassette into the 3’ gRNA target sequence. As the 3’ gRNA target sequence and PAM site are part of coding DNA for last 9 amino acids in exon 6 of *Slc35f6*, we made non-sense mutations to the coding sequence that is 5’ adjacent to in-frame fused APEX2 expression cassette. This mutated sequence does not contain the 3’ gRNA target site and PAM, and therefore is not subject to the Cas9 cleavage, which otherwise only targets the 3’ gRNA target sequence presented downstream of the expression cassette in the *Slc35f6* dRT (Fig. S1b). We also included 40 bp sequences, which are the endogenous DNAs 5’ (upstream) adjacent to the 5’ cleavage site and 3’ (downstream) adjacent to the 3’ cleavage site, to flank the FRT3 upstream and the expression cassette downstream, respectively, to complete EIF construction (Fig. 1b, S1a-c, and S4a). The 40-bp sequences, traditionally referred to as mini-homology-arms (Aida et al., 2016), are exposed upon Cas9 cleavages *in situ* and become DNA end sequences that overlap at the corresponding genomic and EIF ends. We speculated that the short stretch of identical DNA sequences at the corresponding genomic and EIF ends provide defined sequence substrates for DNA repair. Upon 5’→3’ end resection in the initial step of DNA repair, complementary 3’ overhang sequences would be formed in the overlapping DNA sequences, which thereby serves as overhang substrate sequence (OSS). The EIF is further flanked by an identical engineered gRNA target site (Aida et al., 2016) when cloned into a plasmid. Finally, restriction enzyme digestions of the plasmids produce linear dRT fragments with partial vector backbone sequences included at each end to “mask” OSS for protection of EIF ends (Fig. 1b, S2 and S4a).

We predicted that the 40-bp overlapping homology sequences as OSS would be adequate to warrant creations of complementary 3’ overhang sequences in case the ‘end clipping’ (20 bp in mammalian cells) precedes as the initial phase of end resection (Ceccaldi et al., 2016; Truong et al., 2013). Annealing of perfectly or partially complementary 3’ overhangs at the corresponding EIF and genomic ends, followed by either “gap” fill-in DNA synthesis or flap trimming (when necessary) and ligation, results in “paste” of the EIF to fill in the genomic gap created by excision of GTS and ultimately incorporation of altered nucleotide compositions at designed locations (Fig. 1a-1b and S4a).

### Introduction of iCAP designed CRISPR/Cas9 components and dRTs into mouse zygotes and generation of live animals

First, we introduced Cas9 mRNA together with *Slc35f2* dRT and cognate sgRNAs into mouse zygotes by microinjections. To prevent premature dRT cleavage outside of the pronuclei, we did not use a pre-mixed CRISPR/Cas9 ribonucleoprotein complex (RNP), a commonly used practice for introducing CRISPR/Cas9 components into zygotes or cells. We previously observed that RNP cleaves target DNA templates very efficiently at room temperature *in vitro* (data not shown). Nine animals were derived from 54 injected zygotes with 5 mice carrying edited *Slc35f2* alleles as determined by SURVEYOR nuclease assays (data not shown). Genotyping by PCR identified one animal (Animal G) among the 5 mice as a potential heterozygote likely carrying the edited allele designed by iCAP genome editing (Fig. 1c and Table S1).

To evaluate multiplex editing potentials and usage of dRTs at its isogenic genomic locations, we next introduced Cas9 mRNA together with *Slc35f6* and *P2y14* dRTs and cognate sgRNAs into mouse zygotes by co-microinjections. Sixteen animals were derived from 65 injected zygotes. Among these mice, the edited *Slc35f6* alleles are present in 12 mice and the edited *P2y14* alleles are in 10 animals, as determined by SURVEYOR nuclease assays (data not shown). PCR genotyping identified two animals (Animal D and J) and one animal (Animal N) as potential heterozygotes likely carrying iCAP edited *Slc35f6* and *P2y14* alleles respectively (Fig. 1d, S4b and Table S1).

### iCAP enables precise insertion of exogenous sequences in two locations at *Slc35* Loci

We sequenced the genomic region containing the expected edits to determine if the insertion in two locations occurred as designed. In Animal G harboring the edited *Slc35f2* locus, the 5’ end of the EIF was seamlessly pasted to the corresponding genomic end at the 5’ cleavage site in intron 7-8 with FRT3 site inserted exactly as designed (Schematic and *SEQ. I* in Fig. 2a). This observation suggests the complementary 3’ overhangs might have been generated within OSS at the genomic and EIF ends following Cas9 cleavage and facilitated an error-free end joining at the 5’ paste site. Downstream from the 5’ paste site, the sequence remains as that of EIF through to its 3’ end and then transits to an uncut engineered gRNA target site and a section of vector backbone sequence that were included to flank the 3’ end of EIF in the dRT. This indicates that cleavage did not occur at the engineered gRNA site *in situ*. As a result, the OSS at the 3’ end of the EIF was “masked” by the vector sequence that became the new and undefined end. Additionally, sequence at the 3’ paste site also shows a section of 15-bp were deleted from the genomic DSB end at the 3’ cleavage site following Cas9 cleavage. A DNA linker of approximately 2.7 kb was somehow added to bridge the undefined vector sequence retained at the EIF 3’ end and the genomic end for linkage (Schematic and *SEQ. II and III* in Fig. 2a).

**Figure 2.**
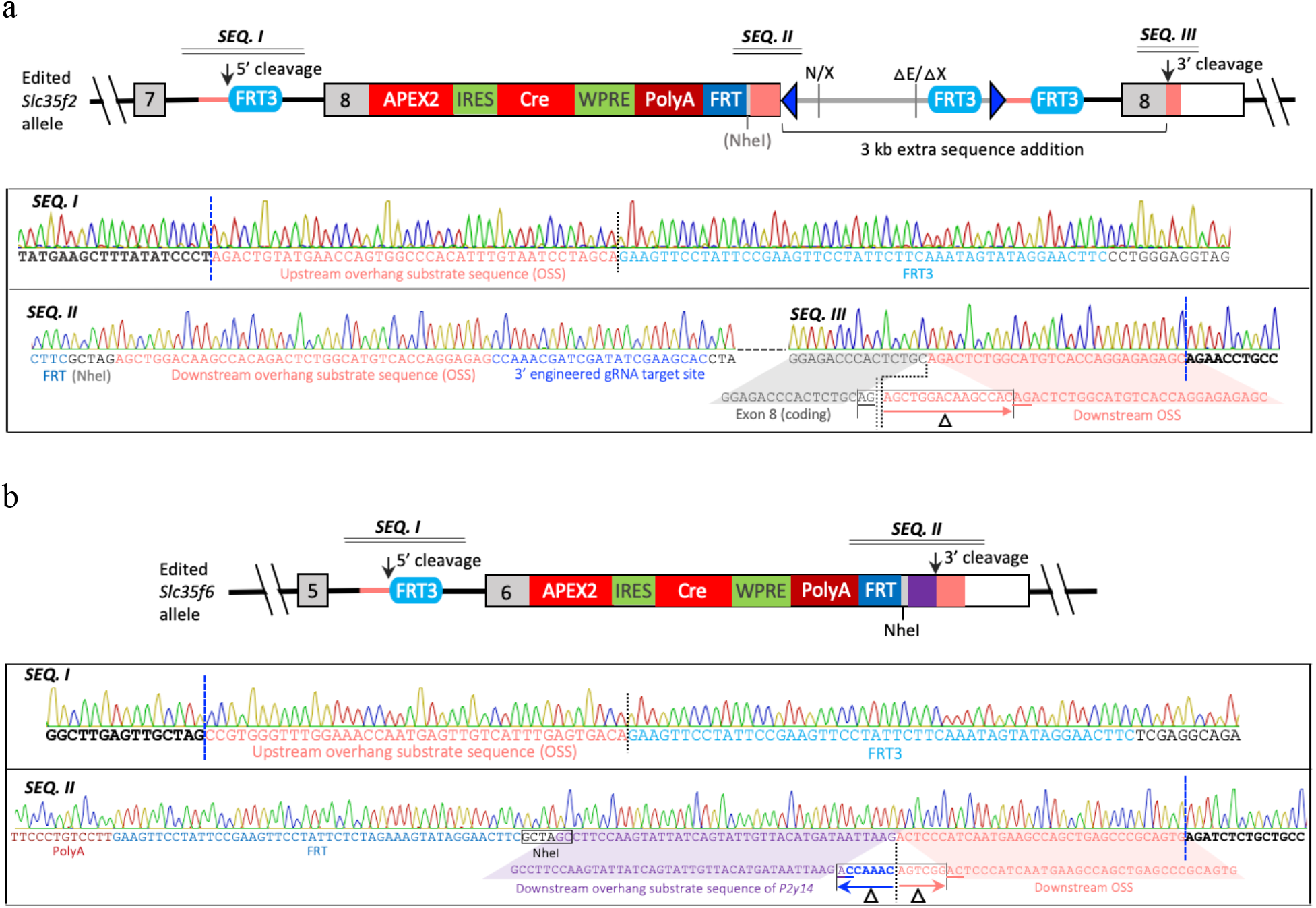
DNA sequence of iCAP edited loci in animals. **(a)** Schematic and sequences of the edited region of *Slc35f2* gene locus in Animal G. The end-joining at 5’ cleavage site is error free. The dRT’s uncut 3’ end (indicated by N/X) and the genomic DSB end at 3’ cleavage site are bridged by a 2.7 kb linker originated from a contaminated vector backbone fragment (N/X-△E/△X) and an uncut 5’ portion of a second copy of dRT (△E/△X-3’cleavage site) resulting in a total of 3.0 kb extension at the 3’ end of EIF (top). ***SEQ. I***, sequence spanning the end-joining at 5’ cleavage site; ***SEQ. II***, sequence region of the EIF’s 3’ end with an uncut engineered gRNA target sequence; ***SEQ. III***, sequence showing the end-joining at 3’ cleavage site. Vertical grey dotted line indicates the linker’s 3’ DSB end generated by possible Cas9 cleavage at a cryptic site near exon8-APEX2 junction in the second copy of dRT molecule. (**b)** Schematic and DNA sequences of iCAP edited *Slc35f6* gene locus in Animal D. The end-joining at 5’ cleavage site is error free. An extra 40-bp sequence (purple box) links the EIF’s 3’end with the genomic DSB end at 3’ cleavage site (top). ***SEQ. I***, sequence showing the end-joining at 5’ cleavage site; ***SEQ. II***, sequence region showing the end-joining at 3’ cleavage site with addition of extra 40-bp sequence (purple). Black bold bases flanking blue vertical dash lines, sequences not included in dRT; vertical black dotted lines, locations of 5’ and 3’ cleavage sites; underlined nucleotide (nt), 2-nt micro-homology ends; delta sign, deletions. Restriction sites: N, NdeI; X, Xmal; E, EagI; grey NheI, the site lost; black NheI, the site retained.

The linker sequence is composed of a truncated EagI-XmaI vector backbone fragment which is likely a contaminant unfortunately co-isolated with the dRT fragment from the construct-containing plasmid (Fig. S2a), and the 5’ portion of a second copy of the dRT. Interestingly, cleavage failure also occurred at the engineered gRNA target site located near the 5’ end of a second dRT molecule in the linker despite a successful cut at the same location in the first copy of the dRT. The observed inconsistency in Cas9 cleavage of the same gRNA target sequence at different locations was also seen at the exon8-APEX2 fusion junction. At this site, a new PAM site was unintentionally constituted within the seed sequence of the original 3’ genomic gRNA target sequence within which the APEX2-IRES-CRE-WPRE cassette was inserted in the dRT (Fig. S1a). It is likely that the repeating 5’ portion of dRT in the linker is a truncated product of Cas9 cleavage near the exon8-APEX2 junction of a dRT molecule. However, this cleavage did not occur at the identical junction sequence in the first copy of the dRT nor was it observed in an *in vitro* cleavage assay to test excision of EIF from the dRT fragment. Nevertheless, the joining of two incompatible DNA ends at the 3’ paste site appears to be facilitated by annealing of two-nucleotide (<u>AG</u>) microhomologies (*SEQ. III* in Fig. 2a).

DNA sequences were also determined for the iCAP edited region of *Slc35f6* locus in Animal D, derived from a zygote that received both of *Slc35f6* and *P2y14* dRTs as a test for dRT usage and multiplex editing. End-joining at the 5’ paste site was also flawless with FRT3 inserted in intron 5-6 as designed (Schematic and *SEQ. I* in Fig. 2b), indicating the likelihood of complementary 3’ overhangs formation within OSS at the corresponding genomic and EIF ends following Cas9 cleavage. Toward the 3’ direction from the 5’ paste site, the sequence remains the same as that of *Slc35f6* EIF until the FRT site at the 3’ end of the expression cassette. The sequence then transits to a fully retained 6-bp NheI restriction site and a 40-bp OSS which was constructed at the 3’ end of *P2y14* EIF. The sequence then continues into the genomic end that lost six nucleotides from the predicted 3’ cleavage site (Schematic and *SEQ. II* in Fig. 2b and Fig. S1b). Sequence structure at the junction of 3’ paste site indicates that OSS at the 3’ end of *Slc35f6* EIF was completely replaced with the 40-bp OSS coming from the 3’ end of *P2y14* EIF. The corresponding genomic and *Slc35f6* EIF ends with unmatched OSS at the downstream genomic cleavage site appear to be linked by annealing of two-nucleotide (<u>AC</u>) microhomology sequence (*SEQ. II* in Fig. 2b).

It is possible that OSS at the 3’ end of *Slc35f6* EIF was exposed following cleavage and subsequently deleted as a result of excessive “end clipping”, resulting in a new and undefined end incompatible to the corresponding genomic end. The available OSS at the 3’ end of the *P2y14* construct may have served as a linker or donor sequence to bridge the incompatible *Slc35f6* EIF and corresponding genomic ends for joining, similar to ssODN bridging (Yoshimi et al, 2016). Alternatively, it can be a result of nonallelic gene conversion (Chen et al., 2007; Harpak et al., 2017) between the two different dRTs as they contain identical expression cassette sequences for most of their length with variations only at the two distal ends (Fig. S3). DNA sequencing of the entire iCAP edited region in the *Slc35f6* locus revealed the correct 5’ portion of the *Slc35f6* EIF with matching isogenic genomic sequence and APEX2-Cre reporter (Fig. 2b). Therefore, the presence of the NheI restriction site and OSS that match to the 3’ end section of *P2y14* EIF strongly indicates gene conversion (Fig. S3 and *SEQ. II* in Fig. 2b). Although the exact pathway for the OSS exchange at this junction is not known, the unrelated 40-bp sequence does not alter precise insertion of the expression cassette as designed for *Slc35f6* locus. Finally, sequence analysis of the edited *Slc35f6* locus in the second mouse (Animal J) revealed it was a partial knockin with only the expression cassette inserted via HR at the targeted location. This phenomenon is often associated with a partial replacement in HR (Becher et al., 2018; Chen et al., 2019).

### Isogenic dRTs guide DNA donors to corresponding locations and recombine with the genome in correct orientation

In order to evaluate precise multiplex editing and usage of dRT to replace GTS by iCAP, we simultaneously co-injected *Slc35f6* and *P2y14* dRTs with cognate sgRNAs and Cas9 mRNA into zygotes. Among sixteen mice produced, no single animal harbored both of precisely edited *Slc35f6* and *P2y14* alleles although many mice carried multiplex edited alleles with indels at the genomic gRNA target sites. However, three heterozygous animals were identified to carry targeted modifications using DNA donors at either *Slc35f6* or *P2y14* loci. As described above, Animal D carrys iCAP edited *Slc35f6* allele while Animal J contains edited *Slc35f6* locus by partial HR.

PCR genotyping identified Animal N as a candidate bearing the iCAP edited *P2y14* allele (Figure S4b). Sequence analysis for the edited region revealed that the EIF was excised from *P2y14* dRT and its 5’ end was joined with the corresponding genomic end with no error (*SEQ. I* in Figure S4c). This flawless end-joining junction at upstream genomic cleavage sites (5’ paste site) was also seen in iCAP edited *Slc35f2* and *Slc35f6* genes. However, the EIF’s 3’ end with OSS was bridged and joined to the 3’ DSB end at the upstream (5’) genomic cleavage site rather than at the downstream (3’) gRNA target site (Figure S4c). The bridge is a 187 bp sequence comprised of a 44 bp vector backbone designed to mask OSS at the 5’ end of *P2y14* dRT and a 143 bp sequence identical to part of the mitochondrial cytochrome oxidase subunit I (*Cox1*) gene. The integration of *P2y14* EIF at a single genomic cleavage site is likely the result of failed cleavage at 3’ genomic gRNA target site. Although Animal N expresses the in-frame fused gene cassette (data not shown), we do not count such edited *P2y14* as an iCAP edited allele. It is worth to note that none of the 10 mice identified to carry edited *P2y14* alleles show edits (indels) at this downstream target site (data not shown), indicating that the cognate sgRNA failed to target Cas9 cleavage at the site *in situ*. Nevertheless, correct usage of dRTs in the right orientation at each edited allele suggests that isogenic genomic sequence in dRTs helped to guide donor DNA to the right genomic locus when two different dRTs are present in a nucleus.

### iCAP edited *Slc35f2* and *Slc35f6* alleles are heritable and expressed properly

The primary goal of editing the *Slc35* gene loci was to generate an in-frame fusion with an APEX2-IRES-CRE-WPRE expression cassette. Successful reporter gene construction requires both germline transmission and faithful expression of fused genes. To evaluate the heritability of iCAP edited alleles, heterozygous male founder Animals G and D were bred with wild-type B6 female mice to derive F1 progeny. Genotypic analysis of the offspring identified F1 animals carrying iCAP edited *Slc35f2* or *Slc35f6* alleles, confirming germline transmission of the edits.

Gene expression from iCAP edited *Slc35* alleles in F1 heterozygotes was also investigated by RT-PCR in brain and kidney tissues where the endogenous genes are highly expressed. Both organs exhibit robust mRNA expression from the edited *Slc35* alleles (Fig. 3a and 3b) with levels similar to those transcribed from wild-type alleles. Sequence analysis of the RT-PCR products revealed proper RNA splicing to remove intron 7-8 in *Slc35f2* and intron 5-6 in *Slc35f6*, indicating no interference by FRT3 insertions, and correct sequences at in-frame fusion junctions between the last *Slc35* codons and the first APEX2 codon (sequence panels in Fig. 3a and 3b). These results demonstrate that the iCAP edited alleles and the in-frame inserted APEX2-IRES-CRE-WPRE expression cassettes are properly expressed in vivo.

**Figure 3.**
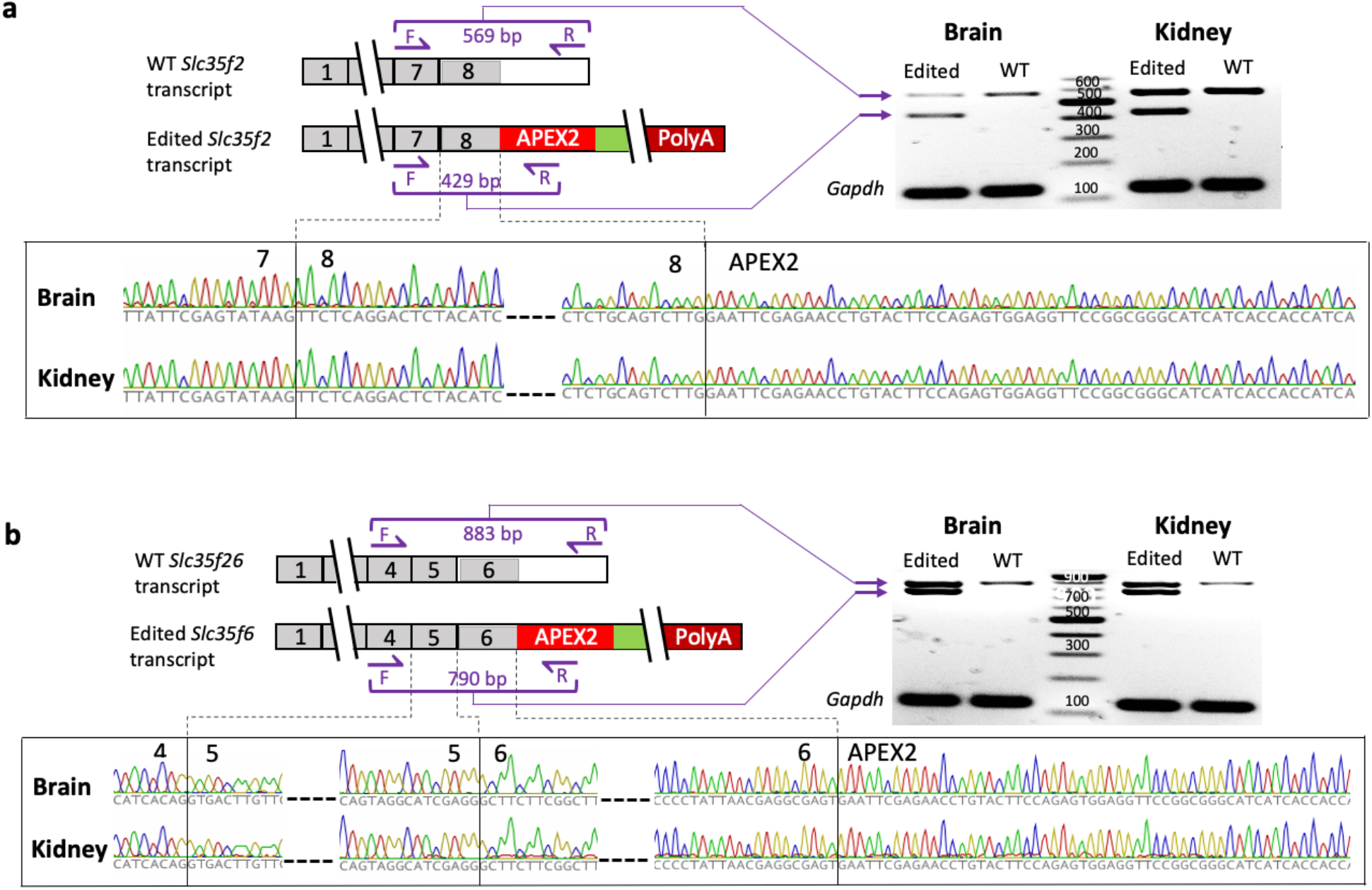
Expression of transcripts from iCAP edited mouse alleles. **(a)** Schematic of transcripts expressed from WT and edited *Slc35f2* alleles (top left). Purple arrow, location of primers used for RT-PCR. Agarose gel image showing specific RT-PCR fragments corresponding to the wildtype and edited *Slc35f2* transcripts expressed in tissues of brain and kidney where the gene is highly expressed (top right). Sequences of RT-PCR fragments showing the correctly spliced junction of exon 7-8 and in-frame expressed reporter gene APEX2 at the boundary (exon8-APEX2) between the last codon of *Slc35f2* and the first codon of APEX2 (bottom). **(b)** Schematic showing transcripts expressed from WT and edited *Slc35f6* alleles (top left). Purple arrow, location of primers used for RT-PCR. Agarose gel image showing specific RT-PCR fragments corresponding to the wildtype and edited *Slc35f6* mRNAs in tissues of brain and kidney where the gene is highly expressed (top right). Sequences of RT-PCR fragments showing the correctly spliced junctions of exon 4-5 and 5-6 as well as in-frame expressed reporter gene APEX2 at the boundary (exon6-APEX2) between the last codon of *Slc35f6* and the first codon of APEX2 (bottom). A 123-bp RT-PCR fragment was generated from the endogenously expressed *Gapdh* transcript as a positive control. Lane Edited, tissues of edited mice; Lane WT, tissues of wildtype mice.

### Design of iCAP using Cas9 or Cas12a for human genome editing

Our results in mouse genome editing indicate that iCAP is useful for inserting large DNA cassettes with high fidelity. Next, we wanted to determine the flexibility and efficacy of using iCAP to alter a single nucleotide. *MED13L* haploinsufficiency syndrome, a genetic disorder in children (Asadollahi et al., 2013), is associated with a spectrum of *de nova* heterozygous loss-of-function variants in *MED13L* gene (Adegbola et al., 2015; van Haelst et al., 2015; Cafiero et al., 2015; Asadollahi et al., 2017; Smol et al., 2018). A single thymine duplication at codon 1497 in exon 20 of *MED13L* gene with 31 exons was identified as a causative gene variant in a *MED13L* syndrome patient, resulting in a S1497F variation (*MED13L^S1497F^*) and consequently reading-frame shift. Furthermore, DNA sequence annotation of the gene variant revealed a premature transcription termination in exon 21 causing a deletion of approximately 690 amino acid residues at the C-terminus of MED13L. We decided to eliminate the single nucleotide mutation in exon 20 through an exon exchange by iCAP genome editing using either CRISPR/Cas9 or CRISPR/Cas12a system.

Similar to iCAP design for mouse genome editing, we identified gRNA target sequences for both Cas9 and Cas12a within intronic regions flanking exon 20 (Fig. 4a, 5a and S5a, S6a). We generated a 561-bp genomic DNA fragment containing wild-type exon 20 (193-bp) and flanking intronic sequences (219-bp 5’ of exon 20 and 149-bp 3’ of exon 20) containing the gRNA target sites. We then inserted a puromycin resistant gene expression cassette (Puro) in the 5’ intron with the insertion site at 25-bp upstream of exon 20. Use of the selection marker eliminates cells harboring indels at the Cas effector cleavage sites, a predominant editing outcome (Cong et al., 2013; Zetsche et al., 2015). The selection marker also serves as a unique tag for the iCAP edited *MED13L* allele. The Puro-exon20 sequence is the core section of EIF to be built in dRTs (Fig. 4a, S7b and 5a, S8b).

**Figure 4.**
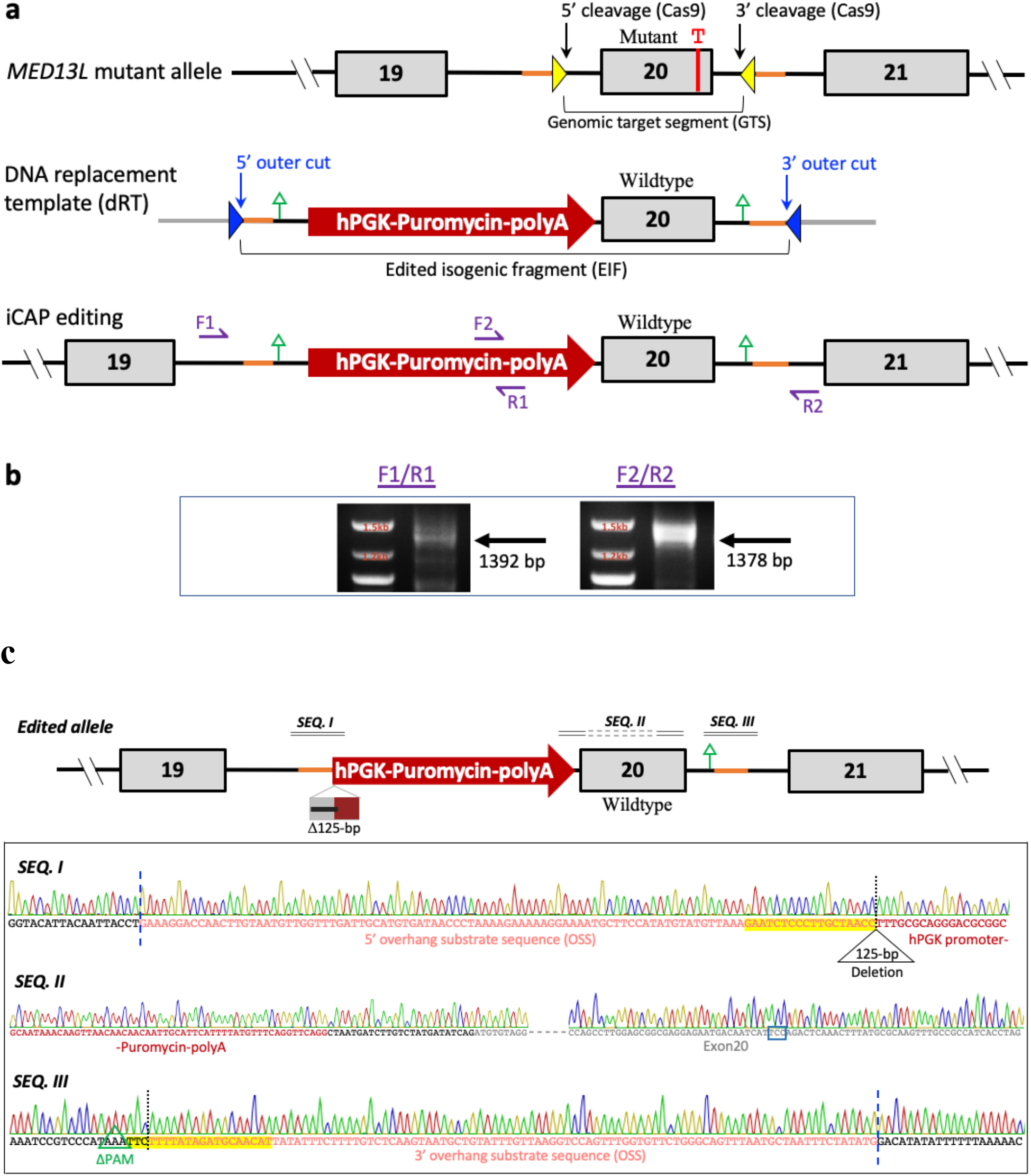
Cas9 mediated iCAP genome editing of *MED13L* locus for exchange of exon 20. **(a)** Schematic of the human *MED13L* locus showing mutant exon 20 and flanking regions in *MED13L^S1497F^* patient cells (top). A thymine duplication is indicated as a T in red, locations of gRNA target sequence as yellow arrowheads and 100-bp overhang substrate sequence (OSS) in salmon lines. Schematic showing the design of a DNA replacement template (middle). Gray lines, parts of vector backbone sequences; blue arrowheads, engineered gRNA target sequence; salmon lines, 100-bp OSS; green vertical arrows, location of mutated PAM sites associated with genomic gRNA target sequences flanking exon 20. Schematic illustrating predicted structure of the edited region as a result of iCAP replacement of GTS with the EIF excised from dRT (bottom). Purple F1/R1 and F2/R2, location of primers for genotyping. (**b)** Genotypic identification of iCAP edited *MED13L* allele in CRISPR/Cas9/dRT transfected patient cell populations. Arrows indicate expected PCR products specifically amplified from iCAP edited alleles. **(c)** DNA sequence of iCAP edited *MED13L* allele. Schematic of the edited region confirmed as a result of replacement of GTS with the EIF (top). Δ, deletion; green vertical arrow, location of the mutated PAM site in EIF. Sequences spanning junctions at 5’ and 3’paste sites and codon 1497 (bottom). ***SEQ. I***, junction at 5’ paste site showing loss of 55-bp intronic DNA and first 70-bp hPGK promoter sequence of the Puro-cassette. ***SEQ. II,*** sequence of Puro cassette polyA and 3’ portion of exon 20 showing wildtype codon 1497 in blue frame. ***SEQ. III,*** error-free end joining at the junction of 3’ paste site. Bases 5’ and 3’ of a vertical blue dash line, sequence not included in the dRT; vertical black dotted lines, the 5’ and 3’ cleavage sites; yellow shaded bases, parts of gRNA target sequences retained; Bases in black, intron sequence; bases in grey, exon 20 sequence.

**Figure 5.**
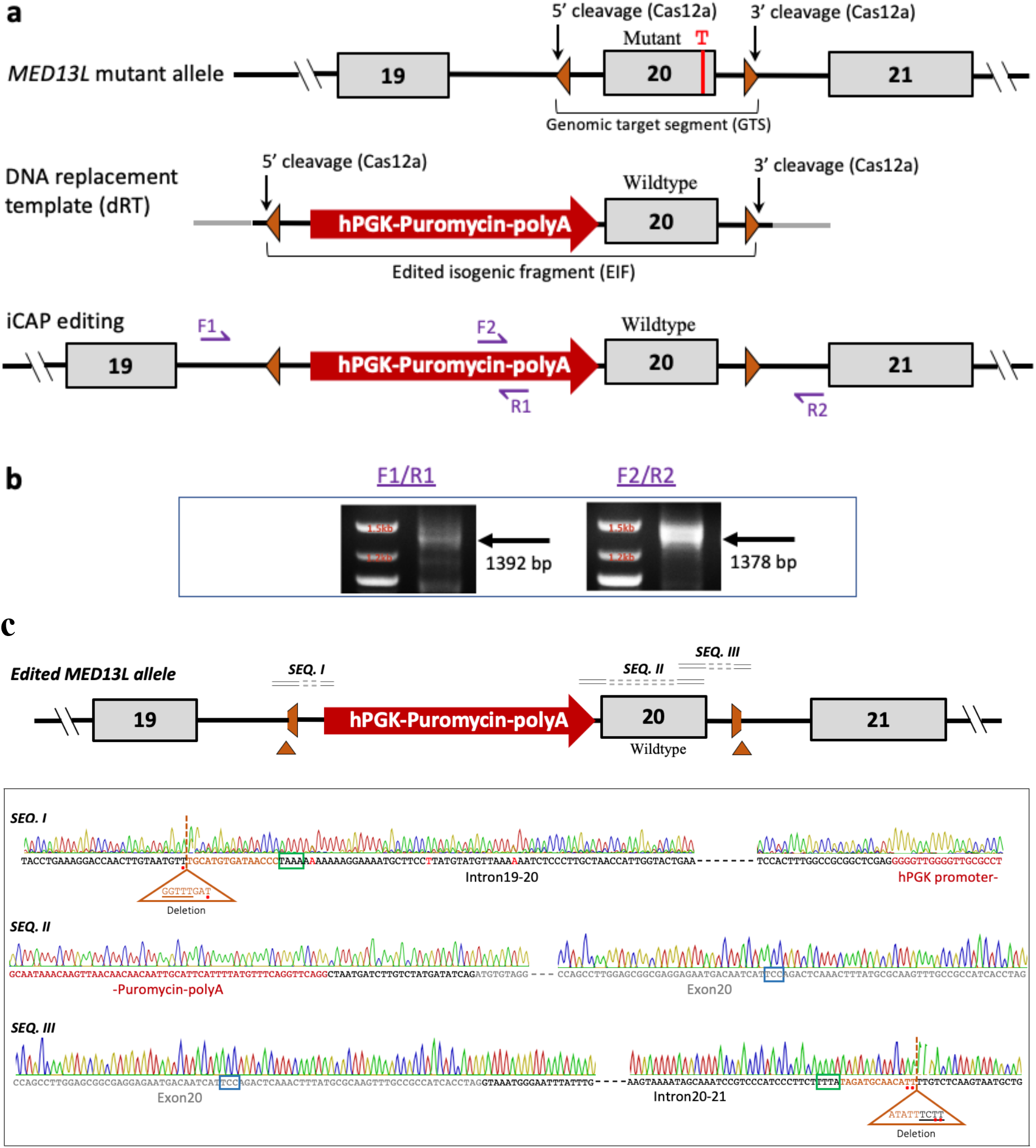
Cas12a mediated iCAP genome editing of *MED13L* locus for exchange of exon 20. **(a)** Schematic of the human *MED13L* locus showing mutant exon 20 and flanking regions in *MED13L^S1497F^* patient cells (top). A thymine duplication is indicated as a T in red, locations of gRNA target sequence as brown arrowheads. Schematic showing the design of a DNA replacement template (middle). Gray lines, parts of vector backbone sequences. Schematic illustrating predicted structure of the edited region as a result of iCAP replacement of GTS with the EIF excised from dRT (bottom). Purple F1/R1 and F2/R2, location of primers for genotyping. (**b)** Genotypic identification of iCAP edited *MED13L* allele in CRISPR/Cas12a/dRT transfected patient cell populations. Arrows indicate expected PCR products specifically amplified from iCAP edited alleles. **(c)** DNA sequence of iCAP edited *MED13L* allele. Schematic of the edited region confirmed as a result of replacement of GTS with the EIF (top). Brown triangle boxes indicate parts of gRNA target sequences deleted. Sequences spanning junctions at 5’ and 3’paste sites and codon 1497 (bottom). ***SEQ. I***, junction sequence at 5’ paste site showing deletion of 8-bp from the upstream (5’) gRNA target sequence in the 5’ end of EIF. ***SEQ. II,*** sequence of Puro cassette polyA and 3’ portion of exon 20 showing wildtype codon 1497 in blue frame. ***SEQ. III,*** junction sequence at 3’ paste site revealing deletion of 9-bp from the downstream (3’) gRNA target sequence in the 3’ end of EIF. Bases 5’ and 3’ of a vertical brown dash line, sequence not included in the dRT; vertical brown dash lines, the 5’ and 3’ cleavage sites; bases in brown, parts of gRNA target sequences retained; bases indicated by red asterisks, possible complementary overhangs of one or two nucleotides (nt) or microhomology nt; underlined bases in deletion boxes, overhang nt generated by Cas12 cut; bases in green frame, PAM sites.

The dRT used for Cas9 mediated iCAP (Cas9 dRT) was built as described above except: i) two genomic gRNA target sites presented in the isogenic genomic segment of EIF were disabled by mutating their cognate PAM sites; ii) the Puro expression cassette was placed at the 5’ end section of EIF rather than at the 3’ end portion as in the mouse dRTs; and iii) the OSS was extended from 40 bp to 100 bp in *MED13L* Cas9 dRT (Fig. 4a and S5b). The increased OSS length may prevent a total loss of the defined ends due to excessive end clipping. Construction of the dRT used for Cas12a mediated iCAP (Cas12a dRT) is relatively straightforward as neither OSS nor artificial engineered gRNA needs to be considered. While Cas9 induces blunt ended DSBs, Cas12a generates staggered DSB with 5’ overhangs of 4 or 5 nucleotides (nt) (Zetsche et al., 2015). We reasoned that we could use Cas12a as a cohesive-end-generating restriction enzyme to excise both GTS from genome and EIF from dRT by targeting the same upstream (5’) and downstream (3’) gRNA target sites flanking GTS and EIF, respectively. The 5’ overhangs of 4- or 5-nt generated at the genomic and corresponding EIF ends are perfectly complementary and could be perceived as defined ends by DNA repair machinery resulting in the joining dependence simplified with the least possibilities of changing the end sequence structures at the junction (Lieber, 2010). Based on this rationale, the Cas12a dRT was built to only contain a fragment that includes the 561 bp genomic DNA sequence and Puro inserted in the intron upstream of exon 20 as described above (Fig. 5a and S6b). The Puro-exon 20 containing fragment is flanked with short vector backbone sequences and become dRTs when isolated from plasmids by SphI-SacI restriction digestions (Fig. S7b and S8b).

### iCAP mediated by Cas9 enables a swap of mutant *MED13L* Exon 20 for WT Counterpart

We introduced the plasmid expressing Cas9 and three sgRNAs (to target 5’ and 3’ cleavage sites on the genome and engineered gRNA target sites on the dRT) together with a 2145-bp SphI-SacI Cas9 dRT fragment (Fig. 4a and S7) into *MED13L^S1497F^* patient fibroblasts by electroporation. The puromycin-resistant cell populations were collected and assigned as 4-1-2 for genotypic analysis after 10 days in selection medium. Using primer pairs of F1-R1 and F2-R2 shown in Fig. 4a, PCR products of approximately 1392 bp and 1378 bp were generated as expected (Fig. 4b), indicating the presence of the edited allele as a result of iCAP replacement of the GTS with the EIF containing WT exon 20 and the puromycin resistant gene.

We next sequenced the PCR products to examine the 5’ and 3’ paste sites as well as the codon 1497 area. In contrast to the iCAP edited mouse genes, the error-free end joining occurred at the 3’ paste site between corresponding genomic and EIF ends in most (4 out of 5) of cloned PCR products sequenced. As shown in *SEQ. III* of Fig. 4c, the 3’ endogenous genomic DNA sequence not present in the dRT is seamlessly transited (3’ to 5’ direction) into the overhang substrate sequence followed by the mutated PAM site that are intronic sequences 3’ of exon 20 as constructed in the EIF. Therefore, the end joining was likely a result of annealing of the complementary 3’ overhang sequences that were formed within the OSS shared at the genomic and EIF ends as intended by iCAP.

Among 4 of 5 cloned PCR amplicons sequenced, junction sequences at 5’ paste site show that fully-retained OSS at the 5’ genomic DSB end was extended by a single cytosine base from predicted Cas9 cleavage position (*SEQ. I* in Fig. 4c and S5a). This extra nucleotide might be added by a template-independent synthesis (Lieber, 2010) or from a shifted cutting site one-base toward the cognate PAM site, a not unusual phenomenon for the endonuclease specificity (Chen et al., 2014). Toward 3’ direction from the extra cytosine, the sequence immediately transited to the hPGK promoter region of the Puro cassette, indicating that deletion occurred at the 5’ end of EIF following excision from dRT by Cas9 cleavage *in situ*. A 225 bp sequence including 100-bp OSS, 55-bp section of intronic sequence with mutated-PAM-site and 5’ first 70-bp of the hPGK promoter region was clipped from the 5’ end of the EIF. A loss of 55-bp intronic and 70 bp hPGK promoter sequences at the junction of 5’ paste site however did not compromise the expression of the Puro gene as the cells with iCAP edited allele are puromycin resistant. Sequence at Puro-exon 20 junction and in the following exon 20 shows the correct sequence with a wild-type (5’-TCC-3’) tri-nucleotide at codon 1497 rather than the codon (5’-TTC-3’) that contains a duplicated thymine in the mutant exon (*SEQ. II* in FIG.4c). These results confirm that the elimination of the single thymine duplication is a result of iCAP genome editing.

### iCAP using Cas12a enables an exchange of mutant *MED13L* exon 20 for WT Counterpart

We also transfected *MED13L^S1497F^* patient fibroblasts with the plasmid expressing Cas12a and the two sgRNAs (targeting 5’ and 3’ cleavage sites on both the genome and dRT) together with a 2139-bp SphI-SacI Cas12a dRT fragment (Fig. 5a and S8). Puromycin-resistant cell populations were collected after 10 days in selection medium and assigned as 5-1-2 (Cas12a) for genotypic analysis. PCR genotyping revealed the presence of the edited allele with amplicons in size predicted for iCAP replacement of the GTS with the EIF (Fig. 5b).

Sequence analysis confirmed the presence of the iCAP edited allele. Interestingly, the paste of both EIF ends to their corresponding genomic ends succeeded with minimal deletions thus undermining possible direct annealing of the complementary 5’ overhangs. In a majority of cloned PCR amplicons, an 8-bp sequence was deleted at the 5’ paste junction (in 7 out of 8 PCR clones sequenced) while a 9-nt sequence was removed at the 3’ paste site in 5 out of 8 PCR clones analyzed (*SEQ. I* and *SEQ. III* in Fig. 5c). The deleted sequences are nucleotides at the 3’ ends of gRNA target sequences (the strand of target sequence), where the cleavage sites are located and distal to PAM sites.

*In situ* Cas12a cleavage at the up- and down-stream gRNA target sites generated staggered DSB ends with the 5’ overhung gRNA target sequences retained at ends of excised GTS and EIF fragments and short complementary 5’ overhangs at genomic ends (Fig. S6). Junction sequence at the 5’ paste site suggests that the complementary 5’ overhang of 5-nt (3’-<u>CCAAA</u>-5’) at the genomic end were deleted resulting in a blunt end while the 5’ overhang of 5-nt (underlined) and adjacent 3-bp (5’-<u>GGTTT</u>GAT-3’) were removed from the 5’ gRNA target sequence retained in the EIF leading to a 5’ blunt-ended EIF (*SEQ. I* in Fig. 5c). Therefore, the end joining at the 5’ paste site was a ligation of these two blunted ends. It is also possible that the joining was facilitated by annealing of a one-base (T:A) complementary 3’ overhangs which may have been generated by limited 5’→3’ end resection at the two defined ends. In a striking similarity, junction sequence at the 3’ paste site reveals end-processing in the same pattern as seen at the 5’ paste site. As shown in *SEQ. III* of Fig. 5c, the complementary 5’ overhang of 4-nt (5’-<u>TCTT</u>-3’) was deleted from the genomic end and the 5’ overhang of 4-nt (underlined) plus adjacent 5-bp (3’-TATAA<u>AGAA</u>-5’) were removed from the 3’ gRNA target sequence retained at the 3’ end of the EIF. As a result of the deletions, both the corresponding genomic and EIF staggered DSB ends were converted to blunt ends for joining at 3’ paste site. Alternatively, 3’ overhangs might be created by limited 5’→3’ resection at each DNA end, resulting in annealing of 2-nt (TT:AA) microhomology DNA presented at each of the two ends.

The minimal deletions that occurred at both of the paste sites might be the result of attempts to avoid reconstitution of the original gRNA target sites at the end joining locations for a stable linkage. The pattern and scale of deletions at the end joining junctions described above are the dominant editing event by iCAP. It appears that deletions of less than 10-bp intron sequences have no deleterious impact on the edited *MED13L* allele that contains tagged Puro-exon 20 with a wild-type tri-nucleotide (5’-TCC’3’) at codon 1497 (*SEQ. II* in Fig. 5c), as evidenced by survival of edited cells under puromycin selection, reversal of an aberrant cellular phenotype as described below, and improvements of cellular structures and functions in the iCAP edited patient cells (Chang, et al., 2022).

### Reduction of *MED13L* mutant alleles in iCAP edited *MED13L^S1497F^* patient cells

As *MED13L^S1497F^* patient fibroblasts contain both mutant and WT *MED13L* alleles, the iCAP mediated mutant exon 20 swapping for the WT counterpart built in the *MED13L* EIF can occur to either alleles. Therefore, we examined the total mismatch reduction at location of the thymine duplication in exon 20 by comparing indel rates between un-edited *MED13L^S1497F^* patient fibroblasts and the non-clonal edited patient cell populations that survived puromycin selection.

SURVEYOR assays were performed according to the strategy shown in Fig. 6a. 270-bp PCR fragments that contain codon 1497 of *MED13L* gene were specifically amplified using genomic DNA templates isolated from normal, the unedited and edited patient cells. The resultant fragments digested by SURVEYOR nuclease were quantified for calculation of indel rates (Cong et al., 2013; Ran et al., 2013). In comparison with the unedited patient cells, there are 6.3% and 7.3% decreases in indel rates in 4-1-2 (Cas9) and 5-1-2 (Cas12a) edited cell groups, respectively. The rate change reflects a mismatch reduction of 20.9% in the 4-1-2 (Cas9) cell group and 24.2% in 5-1-2 (Cas12a) cell populations (Fig. 6b). The mismatch reductions are notably manifested by increased intensities of undigested 270-bp fragment in 4-1-2 (Cas9) and 5-1-2 (Cas12a) cell groups (Fig. 6c). Mismatch reduction calculations can be found in Methods and Table S2. The results indirectly reflect that a single thymine duplication in the mutant allele was corrected in the similar proportions of non-clonal edited cell populations. The estimated efficiency in mutation correction are consistent with the observation of a pathogenic phenotype reversal in which a small but significant reduction in endogenous reactive oxygen species (ROS) levels was detected in non-clonal edited patient cell groups using dihydroethidium (DHE) dye that fluoresces following oxidation (Fig. 6d).

**Figure 6.**
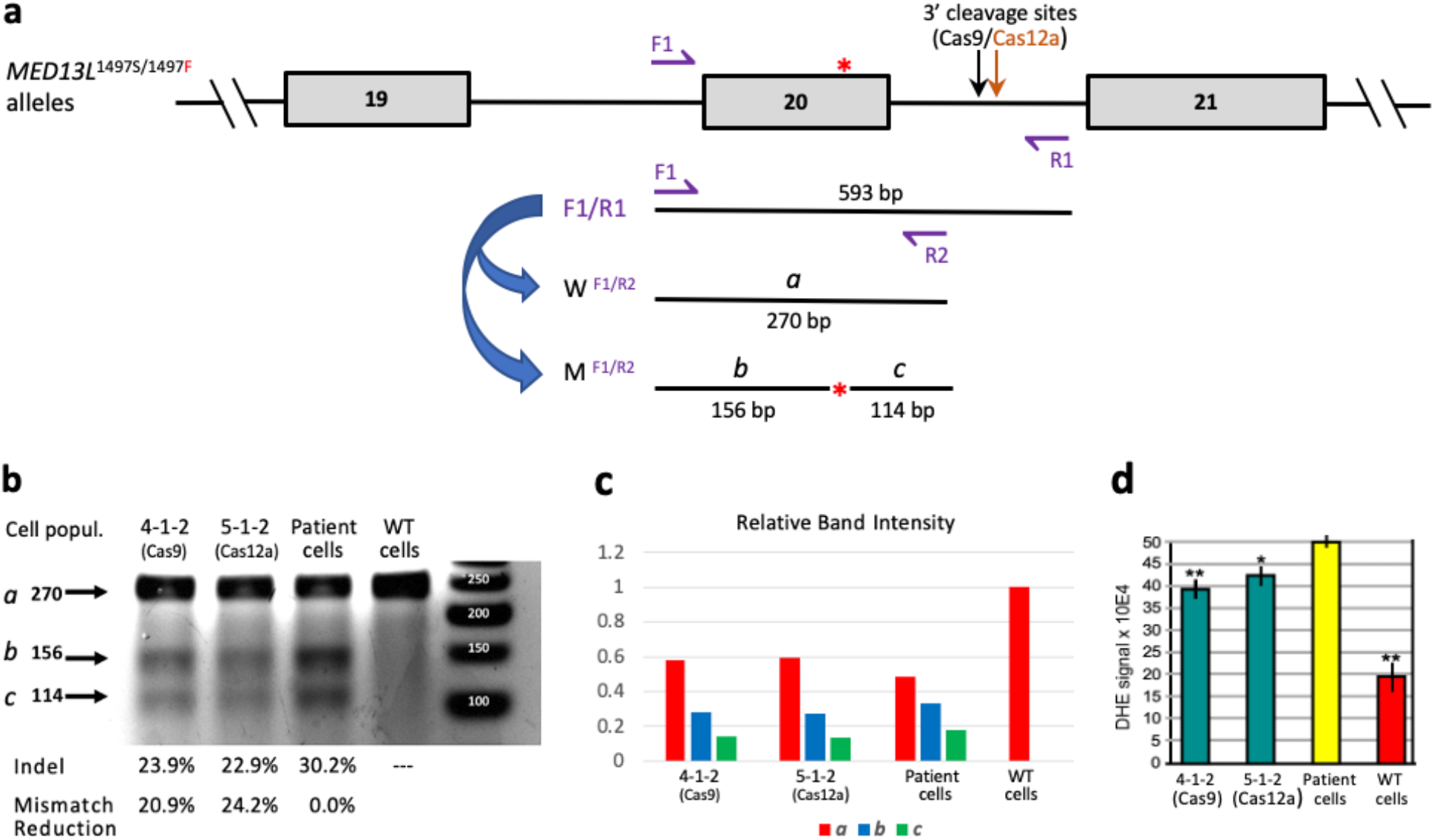
Reduction of mutant *MED13L* alleles correlates to reversal of a cellularly pathogenic phenotype. SURVEYOR assay for evaluation of mismatch reduction at codon 1497 in puromycin resistant *MED13L^S1497F^* cell populations transfected with editing constructs. **(a)** Schematic showing preparation of a specific PCR product for the SURVEYOR assay. The 270-bp product amplified in F1/R2 primed 2^nd^-round PCR contains codon 1497 in exon20 as a sole mismatch location indicated by the red asterisk**. (b)** SURVEYOR assay for mismatch at codon 1497 in un-edited *MED13L^S1497F^* patient fibroblasts (Patient cells), the non-clonal edited patient cell populations transfected with either dRT/sgRNA/Cas9 (4-1-2) or dRT/sgRNA/Cas12a (5-1-2), and WI-38 normal cells (WT cells). Determinations of indel and mismatch reduction can be found in Table S2. **(c)** Quantification of relative band intensities of undigested PCR product *a* and cleavage products *b* and *c* in each group of cell populations. (**d**) Levels of cellular oxidative stress in group 4-1-2 and 5-1-2 of non-clonal edited patient cells (green bars), un-edited *MED13L^S1497F^* patient cells (yellow bar), and wildtype cells (red bar), n=3, ^★^p≤0.05, ^★★^p≤0.01.

### iCAP enables deletion of GTS and end joining of two distant genomic DSB ends

While screening for iCAP edited alleles in animals, we also identified heterozygous mice carrying alleles with “perfectly deleted” GTS. The intervening DNA sequences of 492-bp and 579-bp between predicted CRISPR/Cas9 cutting positions in the 5’ and 3’ gRNA target sequences in loci of *Slc35f2* (Animal A) and *Slc35f6* (Animal J) were precisely deleted, respectively (Fig. 7a-b). The two distant genomic DSB ends generated by Cas9 cleavage were directly re-joined by error-free blunt end ligation, as previously observed in human genome editing (Cong et al., 2013). Therefore, precise deletion of a genomic DNA segment between two gRNA target sites could be another desirable outcome of iCAP genome editing, resulting in a potentially useful by-product as valuable mouse models. Such a perfect deletion also occurred to edited *MED13L^S1497F^* patient fibroblasts that were transfected with the plasmid expressing Cas9 and sgRNAs targeting 5’ and 3’ genomic cleavage sites but no dRT, resulting in deletion of exon 20 from the *MED13L* alleles (Fig. 7c-d) in 12 out of 20 cloned PCR amplicon sequenced.

**Figure 7.**
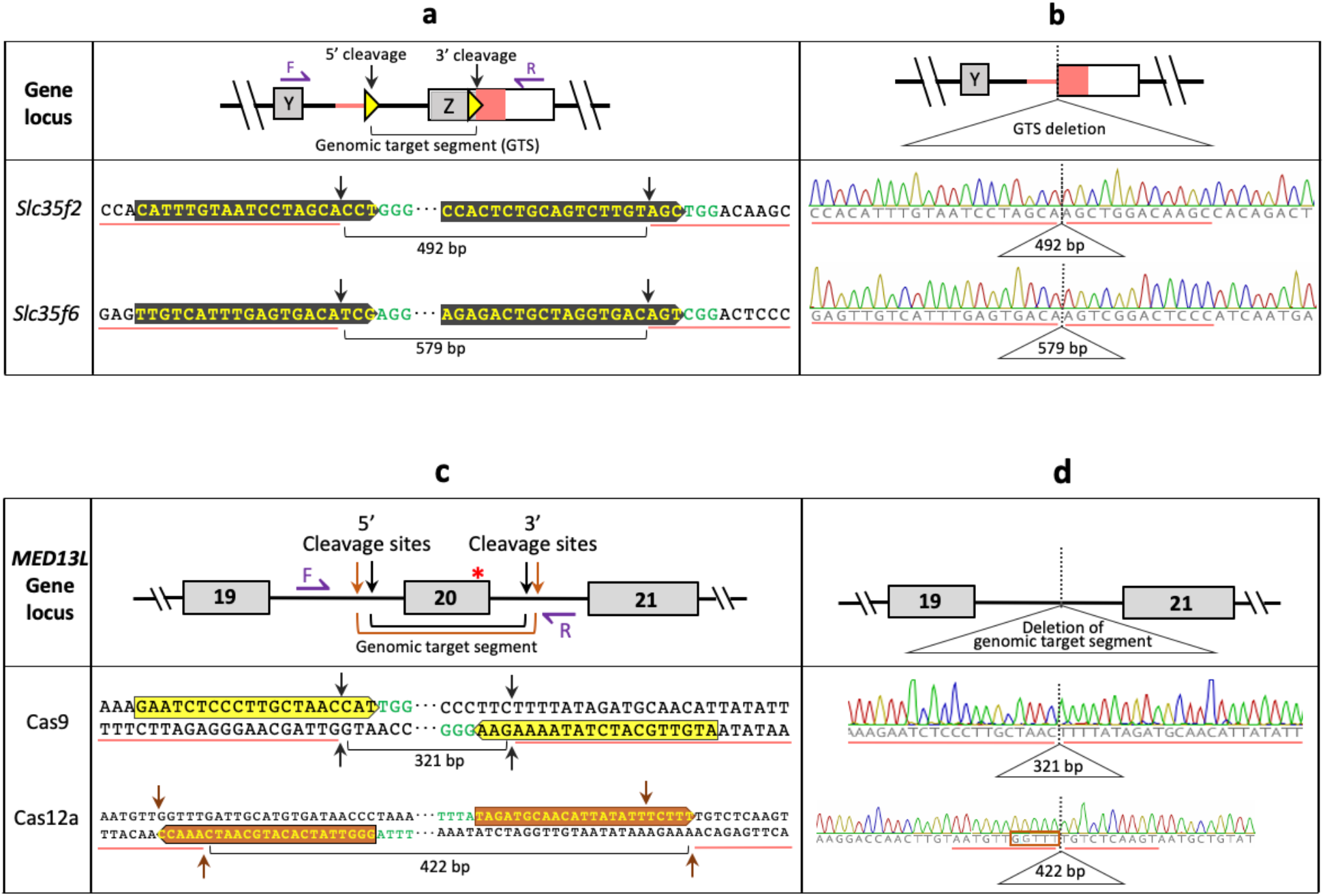
Perfect deletions of genomic target segments and error-free end rejoining without replacement. Schematic of mouse gene loci and sequences at 5’ and 3’ cleavage sites before (**a**) and after (**b**) deletion. Black arrow, locations of DSBs by Cas9 cleavage; yellow arrowhead and bases in yellow, Cas9 gRNA target sequence; bases in green, PAM site; dotted vertical line, boundary of error-free end-joining; underlined bases, genomic sequences flanking cleavage sites; delta frame, deleted GTS. Schematic of *MED13L* allele and sequences at 5’ and 3’ cleavage sites flanking exon 20 before (**c**) and after (**d**) deletion. Black arrow, locations of DSBs by Cas9 cut; orange arrow, locations of DSBs by Cas12a cleavage; red asterisk, location of a thymine duplication; purple arrow, primers for PCR genotyping; bases highlighted in yellow, Cas9 gRNA target sequence; bases highlighted in orange, Cas12a gRNA target sequence; bases in green, PAM site; dotted vertical line, boundary of error-free end-joining; underlined bases, genomic sequences flanking cleavage sites; delta frame, deleted GTS containing exon 20; bases in orange frame, retained 5’ overhang of 5-nt at the staggered genomic DSB end generated by Cas12a cleavage at 5’ cleavage site.

We also performed a transfection experiment in which only the plasmid expressing dual sgRNAs and Cas12a were introduced into the patient cells by electroporation, for *in vivo* evaluations of targeted cleavages at the selected 5’ and 3’ cleavage sites flanking exon 20 of the *MED13L* gene. The cells were cultured for 24 hours after transfection and harvested as pooled cell populations for genotypical screening. PCR genotyping confirmed that the exon20-containing GTS between two gRNA target sites was excised, indicating efficient Cas12a cleavage at the selected sites *in situ* (data not shown).

We further examined how distantly separated genomic ends were joined as short 5’ overhangs generated by Cas12a cut at two different gRNA target sites are not complementary. Sequencing of the genomic region with deleted exon 20 revealed sequences traces at the end-joining junction. A dominant sequence species (14 among 20 cloned PCR amplicon sequenced) shows that the 5’ overhang (3’-<u>CCAAA</u>-5’) at the 5’ genomic DSB end was retained while the 5’ overhang (5’-<u>TCTTT</u>-3’) at the 3’ genomic DSB end was deleted at the end joining (Fig. 7c-d). This junction structure therefore suggests that the two staggered DSB ends with incompatible 5’ overhangs were converted to blunt ends through DNA synthesis at one end and deletion at the other end prior to end joining. Alternatively, the end joining in this particular case could be mediated by direct annealing of partially complementary overhang nucleotides (underlined) at the two staggered genomic DSB ends (3’-CC<u>AAA</u>-5’ at the upstream DSB end and 5’-TC<u>TTT</u>-3’ at the downstream DSB end) with 2-nt (5’-GG-3’) filling-in and 2-nt (5’-TC-3’) trimmed.

Nevertheless, an ‘error-free’ end joining, that occurs between two distant blunt ends created by Cas9 cleavage, can be achieved between two incompatibly staggered DSB ends generated by Cas12a with a ‘micro’ end modification that is limited within the 5’ overhang of 4- or 5-nt. In addition, sequencing traces from the pooled cell populations also suggest the existence of other sequence species showing different patterns of staggered end processing in which the filling-in and deletion of 5’ overhangs occurred in a reciprocal manner as that described above, or either the filling-in or deletion occurred to both ends. We however did not detect any other type of major end modifications at the two distant genomic ends with 5’ overhangs at the end joining boundary, supporting the notion that the DSB ends with 5’ overhangs generated by Cas12a can serve as defined ends that are subject to no or minimal end modifications (Lieber, 2010; Chang et al., 2017).

### No Cas9 and Cas12a off-target activities occur at top off-target sites examined

To evaluate if risks of off-target effects increased with the routine usage of two or three different gRNA target sequences in iCAP genome editing method, we performed targeted amplicon sequencing of 50 off-target sites which present 4 or 5 the most closely mismatched off-target sequences for each of 11 different gRNA target sequences used for the study. We used ICE (Inference of <u>C</u>RISPR <u>E</u>dits) algorithm tool (Conant, et al, 2022) to analyze the sequencing data for evidence of indel mutagenesis at the sites examined. The assessment reports show indel rates of zero percent at these top off-target sites suggesting no off-target effects detected and R^2^ values of 0.95-1.0 indicating “goodness of fit” of the analysis quality (Table S3). Based on the results, iCAP genome editing method does not appear to elevate risks of Cas9 and Cas12a off-target effects.

## DISCUSSION

Precise genome editing via HDR activated by CRISPR/Cas effector induced DSBs often requires a DNA donor of either double-stranded DNA (dsDNA), ssODNs or lssODNs (Cong et al., 2013; Wang et al., 2013; Yang et al., 2013; Yoshimi et al., 2016). The ssODNs (≤ 200 nt) or lssODNs (≤ 2,000 nt) are less suitable as DNA donors for precise editing of longer sequence such as insertion of larger genes, whereas dsDNA templates require homology arms of several-hundred bases to kilobases to be included. When modifications are required at two sites separated by a section of endogenous sequence, HDR with dsDNA donor frequently results in partial knockin of intended edits (Becher et al., 2018; Chen et al., 2019). In contrast, the dRT design strategy for iCAP genome editing is superior to other forms of DNA donors as the DNA template can be as big as 17 kb and constructed using popular pBluescript cloning vectors (Jiang et al., 2002). More significantly, an EIF in dRT can include modifications at multiple sites with no restraint on type of modifications at each site. Additionally, undesired partial knockin can be minimized due to the synchronous excision of the GTS and the EIF to promote *in situ* pasting of the patching fragment to the gapped target locus. Furthermore, gRNA target sequences with the best on-target score and the least off-target effect prediction can be selected as the 5’ and 3’ genomic cleavage sites to flank GTS in iCAP method, overcoming the restraint in selecting a gRNA target site within a 45-bp distance to the modification site in methods using a single genomic gRNA target site for inducing a DSB (Paquet et al., 2016).

An important consideration for placing EIF into the gapped genome is how the junctions of end joining at two paste sites are resolved. The presence of OSS at corresponding EIF and genomic ends facilitated the error-free end joining at the 5’ paste sites at the edited *Slc35f2*, *Slc35f6* and *P2y14* alleles as well as at the 3’ paste site of Cas9 iCAP edited *MED13L* allele. The seamless end joining is likely a result of annealing of complementary 3’ overhangs generated within the OSS on each of the two corresponding DSB ends by 5’→3’ end resection as intended by iCAP. When OSS is absent from the ends of either EIF or genomic DSB, additions of extra DNA sequences at the end joining junction can be a result of DNA repair as seen at 3’ paste sites of Cas9 iCAP edited alleles of *Slc35f2*, *Slc35f6* as well as at the 3’ junction at the paste site of edited *P2y14* allele.

The absence of OSS could be the result of failed *in situ* cleavage at intended target sites resulting in a persistently “masked” OSS. The cause of cleavage failure could be due to the reduced selectivity in guide sequence targeting and/or recognition by CRISPR/Cas9 in *in vitro* vs. *in vivo* contexts (Ran et al., 2015). In addition, sub-optimized amounts of reaction substrates (e.g., sgRNAs, Cas9 mRNA) and/or dRT delivered by microinjections may result in “partial digestion”. Finally, excessive “end clipping” during the initial phase of DNA repair may attribute to OSS loss at the 3’ end of *Slc35f6* EIF and the 5’ end of *MED13L* EIF as observed at the respective end joining junctions.

The distribution of “self” and “foreign” sequence contexts at an end region of EIF may influence how a Cas9-generated DSB is processed by DNA repair machinery. The error-free end joining appears associated with the 5’ sections of *Slc35f2, Slc35f6* and *P2y14* EIFs and the 3’ section of *MED13L* EIF, where DNA is dominantly the endogenous isogenic sequences. Therefore, the sequences in this section appear more like “self” despite having a 48-bp FRT3 exogenous-sequence inserted near the 5’ end of each EIF with isogenic mouse DNA sequence or containing a single nucleotide alteration in the 3’ part of EIF with isogenic DNA sequence of the human gene. In contrast, the end joining with extra sequence addition or deletion at the junctions appears correlated with the 3’ section of *Slc35f6* EIF and the 5’ section of *MED13L* EIF, where DNAs are either a 3.7-kb exogenous expression cassette or 1.5-kb Puro expression cassette that account for 83% and 79% of the entire *Slc35f6* EIF and *MED13L* EIF sequences, respectively. It is possible that the identity of exogenous (“foreign”) sequences renders the DSB end next to them prone to an excessive amount of end processing in search of compatible sequences such as micro-homology nucleotides (underlined in *SEQ. II* of Fig. 2b). These observations may indicate that sequence contexts of “self” or “foreign” identities at DSB ends of EIF may exert influences on the choice of DNA damage repair pathways or the order of actions and iterative processing in a repair pathway (Chang et al., 2017; Lieber, 2010).

Interestingly, “self” and “foreign” sequence contexts at an end region of EIF appear to have no impact on how a Cas12a-generated DSB end is processed. While Cas9 induces blunt ended DSBs, Cas12a generates staggered DSBs with 5’ overhangs of 4 or 5 nucleotides (Zetsche et al., 2015) which may act as a defined end and require fewer NHEJ proteins to prepare the ends for end joining leading to the least variation of the junction resolution (Lieber, 2010). Sequence structures at both 5’ and 3’ paste sites on Cas12a iCAP edited *MED13L* alleles support such a notion as each Cas12a generated DSB end underwent a <10 bp deletion at each of two end joining junctions. The deleted bases are nucleotides at the 3’ end of gRNA target sequences (target sequence strand), where the cleavage sites and 5’ overhang nucleotides reside. The minimal deletions might be an outcome of simplified end processing mediated by the classic non-homologous end joining (c-NHEJ) DNA repair pathway (Chang et al., 2017; Lieber, 2010), resulting in a 5’ overhang DNA end converted to a blunt one. Alternatively, the deletions might be mediated by the alternative-NHEJ (alt-NHEJ) pathway, one of the end-resection homology directed repair pathways (Truong et al., 2013; Lieber, 2010), that scrutinizes for micro-homology nucleotides by clipping DNA ends. The minimal deletions might have arisen from attempts to avoid reconstitutions of the original gRNA target sites, as staggered DSB ends at corresponding genomic and EIF ends are complementary. The deletion of fewer than 10 nucleotides at both paste sites located in introns appear to be tolerated, harmless and manageable as the ‘scars’ do not compromise functions of the iCAP edited allele. Thus, iCAP genome editing using Cas12a is a relatively straightforward approach as neither OSS nor artificial engineered gRNA need to be considered in dRT construction.

Rendering the EIF’s two DSB ends to be detected as “damaged genomic ends” by cellular DNA repair machinery is anticipated for iCAP genome editing. Manifestations of sequence structure at two end-joining junctions indicate that the “damaged” EIF ends are indeed scrutinized and processed by inherent DNA repair pathways for compatibility with corresponding genomic DSB ends. Notably, the end joining at 5’ and 3’ paste sites could be independently mediated by different DNA repair pathways according to the sequence contexts presented at each end section of the EIF to warrant a linkage with genomic ends. Therefore, the operative mode of the iCAP platform can mobilize and harness the utility of multiple DNA repair pathways to “paste” the EIF at two distant DSB genomic ends and patch the genomic gap. This approach circumvents temporal constraints that restrict a particular DNA repair pathway to operate in certain phases of a cell cycle (Ceccaldi et al., 2016) or rely on HR for precise genome editing (Yamamoto and Gerbi, 2018).

Furthermore, the usage of two different gRNA sequences flanking GTS in iCAP genome editing method does not increase risks of Cas9 or Cas12a activities at the top off-target sites as no off-target effects were detected. This is in line with reported findings from studies examining off-target effects in Cas9 involved editing of mouse genome (Ayabe et al., 2019; Hay et al., 2017) and in Cas12a mediated human genome editing (Kim et al., 2016; Kleinstiver et al., 2016).

As a proof of concept, we demonstrate that iCAP is a feasible genome editing platform that enabled versatile and flexible genome editing resulted in a single nucleotide alteration, precisely multi-kilobase DNA insertion and site-specific modifications at multiple locations within a segment of genome. The method can be mediated by either Cas9 or Cas12a or potentially other Cas effectors that induce DSB, thereby significantly increasing availability and choices of gRNA target sites to edit complex mammalian genomes at any positions. The iCAP approach simply requires mediation by orthodox Cas effectors to fulfill multiple types of precise sequence alterations such as insertion, deletion and substitution. Furthermore, iCAP genome editing using Cas12a requires neither OSS nor artificial engineered gRNA target sequences to be considered, significantly simplifying dRT construction. Notably, we demonstrate that an end joining between two distantly and incompatibly staggered DSB ends generated by Cas12a cleavage can be achieved with a minimal end modification confined to 5’ overhang of 4- or 5-nt. Taken together, the demonstrated editing flexibilities make iCAP genome editing highly attractive as a single platform for multiple types of DNA sequence alterations as well as concurrent editing at multiple sites in mammalian genomes.

## MATERIALS AND METHODS

### Analysis of gene loci, editing locations and identifications of gRNA target sites

Genomic DNA sequences of mouse *Slc35f2*, *Slc35f6*, *P2y14* and human *MED13L* were obtained from online resources, (http://uswest.ensembl.org/index.html and https://www.ncbi.nlm.nih.gov), and then catalogued & annotated using Snapgene software (Version 2.1.1) for further sequence analysis. For the mouse genes, sequences of 500 nucleotides, flanking and spanning two editing locations with one at the introns 5’ of the last exon for insertion of a 48-bp FRT3 and the other as proximal as possible to stop codons in the last exon for in-frame insertion of a 3.7-kb expression cassette, were selected and analyzed for identifying gRNA target sites as 5’ cleavage and 3’ cleavage sites recognized by Cas9 endonuclease using the online tool “Benchling” (https://www.benchling.com). For the human gene, intronic sequences flanking exon 20 were analyzed to search for potential gRNA target sequences recognized by Cas9 or Cas12a (Cpf1) programmable nucleases using the same online tool. Based on a category of programmable nuclease (Cas9, Cas12a, etc.), Benchling software analyzed the selected sequences for all available target sites and scores the sites according to established parameters and variables effecting on-target & off-target efficiencies (Doench et al., 2014; Hsu et al.,2013). The gRNA target sites with favorable scores were chosen for validation assay *in vitro*, according to (1) a favorable combination of on-target and off-target scores (an on-target score > 60 and an off-target score close to 100 are optimum) and (2) the close proximity to the desired site of insertion of any exogenous DNA sequence (for example, the endogenous STOP codon of mouse genes, if creation of an in-frame protein fusion is the goal.) or the close proximity to exon20 of the *MED13L* gene from both upstream and downstream directions.

### Cell lines

The patient fibroblast cell line of *MED13L^S1497F^* was obtained following informed consent with the identification of the MED13L Syndrome donor blinded from the participants in this study. The human cell line of WI-38 is normal fetal lung fibroblasts obtained from ATCC (CCL-75, Manassas, VA).

### Confirmation of a thymine duplication in exon 20 of the MED13L mutant gene allele in MED13L Syndrome patient cells

Genomic DNA was extracted from human cell lines of WI-38 and *MED13L^S1497F^* using DNeasy Blood & Tissue Kit (Qiagen, Hilden, Germany) according to manufacturer’s protocols. Each of the genomic DNA samples were used as templates to produce a DNA fragment by PCR with primers pJ327 and pJ328 (Table S4) which are intronic sequences flanking exon 20 of the *MED13L* gene. All primers and oligos in this study were custom-made (sigmaaldrich.com, St. Louis, MO) and listed in Table S4. The PCR primed with pJ327-pJ328 was performed using Phusion High Fidelity DNA Polymerase (NEB, Ipswich, MA) according to the manufacturer’s protocol on an Applied Biosystems 2720 Thermal Cycler. In brief, 25 µl reactions (5 µl 5X Phusion HF/GC Buffer, 2 µl 2.5mM dNTPs, 1.25 µl 10 uM Forward Primer, 1.25 µl 10uM Reverse Primer, various amounts of Template DNA, 0.5 µl Phusion DNA Polymerase, dH2O) were run on a thermal cycler with typical PCR parameters (initial denaturation at 98°C, 3min; 25-35 cycles of 98°C, 15 sec, 45-72°C, 30 sec; 72°C, 30 sec/kb; final extension at 72°C, 7min; 4°C hold). PCR products of approximately 561bp were generated from genomic DNAs of both WI-38 cells and *MED13L^S1497F^* cells, respectively. The products were first incubated with Choice-Taq DNA polymerase (Thomas Scientific, Swedesboro, NJ) at 72°C for 20min and then TA cloned into the <u>pGEM-T Easy</u> vector (Promega, Madison, WI). The cloned PCR products were Sanger Sequenced with M13 Forward (5’-TGTAAAACGACGGCCAGT-3’) and Reverse (5’-CAGGAAACAGCTATGAC-3’) primers and the DNA sequence reads were analyzed with Geneious software (geneious.com, Auckland, New Zealand). A single thymine duplication at codon 1497 in exon 20 was revealed in patient cells, in comparison with the sequence from human WI-38 normal cells, confirming the presence of a mutant *MED13L* allele in the patient cells.

### In vitro transcription of sgRNAs and validation of gRNA target sites

Templates for *in vitro* transcription of SpCas9 guides were amplified from the plasmid pX330 (Addgene, Watertown, MA) using primer pairs with 65 oligonucleotides as a 5’ primer (forward) and the oligonucleotides 5’-AAAAGCACCGACTCGGTGCC-3’ as a 3’ primer (reverse). The 65 oligonucleotides are comprised of 5’-GCGCGCTAATACGACTCACTATAGGNNNNNNNN NNNNNNNNNNNN GTTTTAGAGCTAGAAATAGC-3’, in which NNN- - - represent 20-nucleotide protospacer preceded with a T7 minimal promoter. The protospacer corresponds to genomic gRNA target sequences identified in the sequence regions of *Slc35f2, Slc35f6*, *P2y14* and *MED13L* loci for 5’ and 3’ cleavages on genome and to the engineered gRNA target sequence 5’- GTGCTTCGATATCGATCGTT-3’ (Aida et al., 2016) for outer cuts (relative to positions of the genomic 5’ and 3’ cleavage sites) on dRT. Approximately 120 bp PCR products were produced with various forward primers and the reverse primer pJ161, and each of the PCR products were used as a template for in vitro transcription of a Cas9 sgRNA containing the cognate gRNA target site. Phusion DNA Polymerase (NEB, Ipswich, MA) was used for the amplification according to manufacturer’s protocols.

For generation of DNA templates used for in vitro transcription of Cas12a sgRNAs, a minimal T7 promoter-RNAfold (direct repeat sequence) oligomer pJ227 (<u>GCGCGCTAATACGACTCACTATAGG</u>TAATTTCTACTAAGTGTAGAT), was annealed with variously specific-target-containing crRNA oligomers (pJ343, pJ344, pJ345, pJ346, pJ347, pJ348, pJ362, pJ363, and pJ364) with pJ227 through overlapping direct repeat sequence, followed by extension in a simple PCR reaction to generate approximately 70 bp products. Each of the PCR products were used as a template for in vitro transcription of one Cas12a sgRNA with the cognate gRNA target site in the template.

PCR products produced as templates for in vitro transcription were purified with either the Qiaquick PCR Purification Kit (Qiagen, Hilden, Germany) for Cas9 templates, or with the QIAEX II Gel Extraction Kit (Qiagen, Hilden, Germany) for Cas12a templates, and all templates were treated with Proteinase K (0.2 mg/ml with 0.5% SDS) at 50°C for 30minutes to remove any traces of contaminating RNAse. The *in vitro* transcription was performed using MEGAshortscript T7 Transcription Kit (LifeTechnologies, Carlsbad, CA) following manufacturer’s protocols. After incubation for 4 hours at 37°C, samples were treated with DNase I for 15 minutes at 37°C to remove DNA templates. *In vitro* transcribed sgRNAs were purified and eluted with MEGAclear Purification Kit (LifeTechnologies, Carlsbad, CA) for the Cas9 102 nucleotide RNA products or with the mirVana miRNA isolation kit (Ambion, Austin, TX) for the Cas12a 44 nucleotide RNA products, and were stored at −80°C for subsequent uses.

In an *in vitro* assay to validate sgRNAs on-target cleavage induction, 30 nM *in vitro* transcribed sgRNAs, 3 nM RNAse-free DNA fragments containing gRNA target sequences and 30 nM Cas9 or Cas12a protein (NEB, Ipswich, MA) were mixed in reaction tubes as per manufacturer’s protocol. At the end of the reaction, 1 μl RNAse A was added and then incubated for additional 15 minutes at 37°C to degrade sgRNA. DNA fragments in the reaction were purified with QIAquick PCR Purification column (Qiagen, Hilden, Germany) to remove residual protein, and analyzed by an agarose gel. Validated gRNA target sequences with accurate target recognition and efficient cleavage by CRISPR/Cas were selected in pairs to flank the genomic target segment (GTS) as the 5’ and 3’ cleavage sites. For iCAP editing using Cas9, the selected gRNA target sequences that flank the *Slc35f2* GTS are 5’-CATTTGTAATC CTAGCACCT-3’ (sense strand) as 5’ cleavage site and 5’-CCACTCTGC AGTCTTGTAGC-3’ (sense strand) as 3’ cleavage site; the selected gRNA target sequences which flank the *Slc35f6* GTS are 5’- TTGTCATTTGAGTGACATCG-3’ (sense strand) as 5’ cleavage site and 5’- AGAGACTGCTAGGTGACAGT-3’ (sense strand) as 3’ cleavage site; the chosen gRNA target sequences flanking the *P2y14* GTS are 5’-CCCTGCACCACTCGTCATGT-3’ (sense strand) as 5’ cleavage site and 5’-TGATAATACTTGGAAGGGGA-3’ (anti-sense strand) as 3’ cleavage site; the selected gRNA target sequences flanking *MED13L* exon 20 are 5’- GAATCTCCCTTGCTAACCAT-3’ (sense strand) as 5’ cleavage site and 5’- ATGTTGCATCTATAAAAGAA-3’ (anti-sense strand) as 3’ cleavage site (Fig. S1 and S5). For iCAP editing using Cas12a, the selected gRNA target sequences flanking *MED13L* exon 20 are 5’-GGGTTATCACATGCAATCAAACC-3’ (anti-sense strand) as 5’ cleavage site and 5’- TAGATGCAACATTATATTTCTTT-3’ (sense strand) as 3’ cleavage site (Fig. S6).

### Constructions of sgRNAs/Cas9 and sgRNAs/Cas12a expression vectors for MED13L editing

For construction of the sgRNAs/Cas9 expression vector *<u>MED13L</u>*<u>-Exon20-Cas9-gRNA+protein</u> (Fig. S7a), pairs of complementary oligoes corresponding to each crRNA sequence were denatured in STE buffer (10mM tris, 1mM EDTA, 100mM NaCl) at 100°C for 10 minutes and then annealed slowly by cooling to room temperature. The annealed oligo products were directionally cloned into the Addgene Multiplex CRISPR/Cas9 Assembly Kit (Addgene, Watertown, MA) as crRNA genes for sgRNA expression. In addition to a Cas9 gene, the expression vector contains three crRNA genes for expression of three different sgRNAs with two of them targeting the 5’ and 3’ gRNA sites, that flank exon 20 of the *MED13L* gene as described above and are also present in the dRT containing the cognate PAM sites mutated, and the third one targeting an engineered gRNA target site (GTGCTTCGATATCGATCGTT) flanking the EIF in the dRT (Fig. 4a and Fig. S5). To make sgRNAs/Cas12a expression vector *<u>MED13L</u>*<u>-Exon20-Cas12a-gRNA+protein</u> (Fig. S8a), pairs of complementary oligoes corresponding to each crRNA sequence were processed and annealed in the same condition as above. The annealed oligo products were directionally cloned into a modified version of the gRNA+LbCas12a expression plasmid <u>pTE4398</u> (Addgene, Watertown, MA) as crRNA genes for sgRNA expression. In addition to a Cas12a gene, the vector contains two crRNA genes for expressions of corresponding sgRNAs targeting the 5’ and 3’ gRNA target sites that flank exon 20 of the *MED13L* gene as described above and are also included and preserved in the Cas12a dRT (FIG. 5a and Fig. S6). The expressions of crRNA genes are driven by autonomous human U6 promoters and the expressions of nucleases are driven by either the chicken beta-actin promoter for Cas9 or human CMV promoter for Cas12a in each of the expression vectors. All the inserts cloned into the expression plasmids were confirmed correct by DNA sequencing through custom-ordered sequencing service provided by Genewiz (genewiz.com, South Plainfield, NJ).

### Construction of DNA replacement template (dRT) containing edited nucleotide alterations

The intended editing for the mouse genes was to make nucleotide alterations at two locations in a gene locus, (1) to insert a FRT3 site in the intron 5’ of the last exon and (2) in-frame to insert a 3.7 APEX2-IRES-CRE-WPRE (*Slc35f2* and *Slc35f6*) or EGFP-IRES-CRE-WPRE (*P2y14*) expression cassette immediately 3’ of the last codon of the endogenous gene. Based on analyses of *Slc35f, Slc35f6* and *P2y14* loci, the editing locations, identification and validation of gRNA target sites, we designed the edited isogenic fragments (EIF) in pre-constructed DNA replacement templates to be organized in a 5’ to 3’ direction as: upstream (5’) overhang substrate sequence (OSS) of 40-bp intron sequence, FRT3 (exogenous DNA), intron sequence, coding sequence of last exon, expression cassette (APEX2-3XFlag-2xHA-StopCodon-IRES-Cre-NLS-WPRE-hGHpolyA-FRT), and downstream (3’) OSS (40-bp sequence around endogenous stop codons), as shown in Fig. 1b. As segments of endogenous genomic DNA in each EIF contain genomic gRNA target sequences for 5’ and 3’ cleavage, we disrupted these target sequences by insertion of the FRT3 and expression cassette at these two sites respectively, or by making non-sense mutations to alter the target sequence that is a part of coding DNA, to prevent Cas9 cleavage at the locations (Fig. S1b). The EIF is flanked with engineered gRNA target sequences when cloned into a vector as a final product (Fig. S2). The dRTs are produced as a linear fragment by XmaI-NdeI (*Slc35f2* and *Slc35f6*) or SpeI-NotI (*P2y14*) restriction digestion of the vector with partial vector backbone sequences that flank the EIF. The partial vector sequences, which can be efficiently removed from dRT fragments by CRISPR/Cas9 cleavage at engineered gRNA targets in *in vitro* testing, are designed to protect (mask) OSS at the EIF ends during the course to reach to nucleus for the dRT. Cleavage at the engineered gRNA target sites inside the nucleus (*in situ*) removes the partial vector backbone sequences and expose OSS at both ends of the EIF.

To construct dRT for iCAP *MED13L* editing using Cas12a, a 561-bp DNA product was generated by PCR amplification of the WT *MED13L* allele with primers of pJ327-pJ328. The sequence contains a WT exon 20 of the *MED13L* gene and partial 5’ and 3’ flanking intron sequences containing selected gRNA target sequences and was cloned into a <u>pGEM-T Easy</u> vector to generate the plasmid of <u>MED13L ex20 pJ327-328 in pGEM-T Easy</u>. A puromycin resistant gene unit was amplified from plasmid <u>pGL3-U6-sgRNA-PGK-puromycin</u> (Addgene, Watertown, MA) by PCR and the 1473 bp product was then inserted in an EcoRV site in intron 19-20, 25 bp upstream of exon 20, resulting in the plasmid of *<u>MED13L</u>*<u>-Exon20-Cas12a-dRT</u> (Fig. S8b). SphI-SacI restriction digestions of the plasmid generate the 2139 bp dRT fragment for Cas12a mediated iCAP editing (referred to as Cas12a dRT). Cleavages induced by Cas12a at the 5’ and 3’cleavage sites on the dRT excised a 1920 bp EIF in an in vitro assay.

To construct dRT for iCAP *MED13L* editing using Cas9, the *<u>MED13L</u>*<u>-Exon20-Cas12a-dRT</u> plasmid was used as a template for incorporating 100 bp overhang substrate sequences (OSS) in the dRT construct. The OSS are endogenous DNAs 5’ adjacent to the 5’ cleavage site and 3’ adjacent to the 3’ cleavage site respectively (Fig. S5a). By sequential PCR reactions using primer sets of pJ356 with pJ358 and pJ357 with pJ359 (Table. S3), the OSS were incorporated as 5’ and 3’ end sequences of the EIF which was then flanked by uniquely engineered Cas9 gRNA target sites. In addition, the PAM sites associated with the 5’ and 3’ genomic Cas9 gRNA target sites in the dRT were mutated to prevent cleavages at these two target sites on the dRT (Fig. S5b). The constructed 2060 bp fragment was TA cloned into the <u>pGEM-T Easy</u> vector, resulting in the plasmid *<u>MED13L</u>*<u>-Exon20-Cas9-dRT</u> (Fig. S7b). SphI-SacI restriction digestions of the plasmid generate a 2145 bp dRT fragment for Cas9 mediated iCAP editing (referred to as Cas9 dRT). Cas9 cleavages at engineered gRNA target sites excise a 2026 bp EIF from the dRT in an in vitro assay. The entire dRT fragments were sequenced and confirmed to have correct DNA sequences as designed.

### Animal Production

Mice were contractually produced by the transgenic mouse facility at University of Pennsylvania School of Veterinary Medicine. Protocols and procedures for animal production were approved by the Institutional Animal Care and Use Committee (IACUC). Briefly, an RNase-free buffer solution (5 mM Tris and 0.1 mM EDTA in ultra-pure water, PH 7.4) containing sgRNAs (40-60 ng/μl), dRT (1.5-3 ng/μl) and Cas9 mRNA (100 ng/μl, Trilink Biotechnologies, San Diego, CA) was microinjected into pronuclei and cytoplasm of one-cell stage embryos obtained from superovulated B6D2F1 female mice (Jackson Laboratory, Bar Harbor, Maine). Injected embryos were maintained in M16 medium and cultured for at least one hour in a 100% humidified incubator with 5% CO_2_ at 37°C before implantation. A group of 20 injected embryos on average were transferred into oviducts of a pseudopregnant B6CBAF1 female mouse (Jackson Laboratory, Bar Harbor, Maine) for full-term development. Founder (F0) mice carrying the desired edited allele were genotypically identified and bred with C57BL/6J (Jackson Laboratory, Bar Harbor, Maine) to produce F1 progeny which were screened genotypically for heterozygotes to determine germline transmission of the edited allele.

### Cell culture and transfection of dRTs and sgRNA/Cas expression vectors

Human cell lines of WI-38 and *MED13L^S1497F^* were propagated in culture plates with DMEM (Dulbecco’s Modified Eagle Medium, Corning Inc., Corning, NY) containing 10% fetal bovine serum (Denville Scientific, Holliston, MA) and 100 unit/ml penicillin/100 μg/ml streptomycin (Sigma-Aldrich, St. Louis, MO) and maintained in a 37°C humidified incubator supplied with 5% CO2. For transfection, 1.0X10^6^ of patient cells carrying the *MED13L* mutant allele were suspended in 80 μl of Opti-MEM medium (Invitrogen, Carlsbad, CA). The cell suspension was combined with 20 μl of genome editing constructs containing either 6 μg of Cas12a dRT and 6 μg of the matching sgRNA/Cas12a expression plasmid or 6 μg of Cas9 dRT and 6 μg of the matching sgRNA/Cas9 expression plasmid. The combined suspension of 100 μl was electroporated by the NEPA21 Electro-Kinetic Transfection System (Nepa Gene Co., Ltd, Ichikawa-City, Japan) with parameters of 175 voltage, 2 pulses of 5 msec length and 10% decay rate for pouring pulse and manufacturer pre-set parameters for transfection pulse. The cells transfected with Cas12a dRT and sgRNA/Cas12a expression plasmid were assigned as 5-1-2 while the cells transfected with Cas9 dRT and sgRNA/Cas9 expression plasmid were labeled as 4-1-2. Following transfection, the cells were cultured in the same culture medium for 48 hours and then maintained in the medium supplemented with 1 μg/ml of puromycin (Sigma-Aldrich, St. Louis, MO) for 10 days. Non-clonal surviving cell populations were harvested for genotypic analysis to identify the iCAP edited mutant *MED13L* allele. Another set of transfection experiments were performed as *in vivo* evaluations of cleavages at both the 5’ and 3’ cleavage sites that flank exon 20 in *MED13L* alleles. 1.0X10^6^ of *MED13L^S1497F^* patient fibroblasts suspended in 80 μl of Opti-MEM medium were combined with either 20 μl of 10 μg sgRNA/Cas12a expression plasmid or 20 μl of 10 μg sgRNA/Cas9 expression plasmid with no dRT included, followed by electroporation with the same parameters. The transfected cell populations were assigned as iCAP-Cas12a-Eva and iCAP-Cas9-Eva, respectively, cultured for 24 hours and then harvested for genotypic analysis.

### Genotyping and Sequencing

To isolate genomic DNA from animals, biopsies of tail tips were dissolved in 500 μl lysis buffer (50 mM Tris-Cl, pH 8.0, 50 mM EDTA, 100 mM NaCl, 1% SDS, 0.2 mg/ml Proteinase K). The dissolving solution was incubated overnight at 55°C with agitation followed by removal of tissue debris by centrifugation. Genomic DNA in lysis buffer was precipitated with 1/10th volume 3M NaOAc (pH5.2) and 500 μl isopropanol, and DNA precipitates were resuspended in 100-200 μl TE buffer following clean washes with 70% ethanol. To isolate genomic DNA from cultured cells, 50 μl of cell samples were lysed in an equal volume of Proteinase K (1mg/ml) in PBS with incubation on a thermal cycler at 65°C for 1 hour followed by incubating at 95°C for 20 minutes.

For genotypic screening of animals, primer pairs of F1/R1 (pJ153/dn739 for *Slc35f2,* pJ159/dn739 for *Slc35f6,* pJ187/dn162 for *P2y14*) and F2/R2 (dn1202/pJ154 for *Slc35f2*, dn1202/pJ160 for *Slc35f6,* dn1202/pJ138 for *P2y14*) were used for amplifying the 5’ and 3’ portions of the edited region in the iCAP edited allele, respectively (Fig. 1b, S4a and Table S4). The F1 and R2 primer sequences are corresponding to endogenous genomic sequences that are distantly upstream and downstream of the endogenous isogenic sequence included in the dRT and R1 and F2 primer sequences corresponding to the exogenous expression cassette sequence. For genotypic screening of transfected cells, primer pairs of F1/R1 (pJ375/ pJ361) and F2/R2 (pJ360/pJ355) were used for amplifying the 5’ and 3’ portions of the edited region in the iCAP edited *MED13L* allele, respectively (Fig. 4a, 5a and Table S4). All PCRs were performed using Phusion High Fidelity DNA Polymerase (NEB, Ipswich, MA) according to manufacturer’s protocols on an Applied Biosystems 2720 Thermal Cycler. PCR fragments with expected size were purified and TA cloned in the <u>pGEM-T Easy</u> Vector (Promega, Madison, WI) for DNA sequencing. Sanger Sequencing for cloned PCR fragments were performed through sequencing service provided by Genewiz. All the sequence data obtained were analyzed with Geneious software.

### RT-PCR characterization of gene expression from iCAP edited Slc35f2 and Slc35f6 alleles

Brains and kidneys were collected from euthanized founder or F1 animals carrying edited alleles as well as from wildtype mice. Organs were preserved in RNA*later* RNA Stabilization Solution (Invitrogen, Carlsbad, CA) and stored at −80°C according to manufacturer’s protocols. Total RNA was isolated from each organ using PureLink RNA Mini Kit (Invitrogen, Carlsbad, CA), according to manufacturer’s protocols. Briefly, preserved and frozen brains and kidneys were thawed and approximately 30 mg of the tissues was dissected and placed in 0.6ml of lysis buffer supplemented with 1% 2-mercaptoethanol. Tissues in the buffer were disrupted using an RNase-free teflon pestle, followed by passing 5-10 times through an 18-20 gauged needle for homogenization. Total RNA was extracted by spin-column according to manufacturer’s protocols and eluted from the column in 30 μl of RNase-free water with concentrations determined by Nanodrop analysis. The RNAs were either immediately frozen at −80°C or DNase treated with the TURBO DNA-free Kit (Invitrogen, Carlsbad, CA).

The DNase-treated RNA was used for cDNA synthesis utilizing the ProtoScript First Strand cDNA Synthesis Kit (NEB, Ipswich, MA) according to manufacturer’s protocols. For each synthesis reaction, 125 ng of total RNA was used for both reverse transcription positive and reverse transcription negative reactions with d(T)_23_ oligo-dT primer provided in the kit. The mixture of total RNA and d(T)_23_ oligo-dT primer was first denatured at 65°C for 5minutes, followed by adding M-MulV Enzyme & Reaction Mix to the reverse transcription positive reactions only, and then incubated at 42°C for 1 hour followed by incubation at 80°C for 5minutes to inactivate M-MulV Enzyme. Resulting cDNA was then either used for semi-quantitative RT-PCR analysis or stored at −20°C.

To examine functional transcription of in-frame inserted exogenous expression cassette from iCAP edited *Slc35f2* and *Slc35f6* alleles, the strategy for RT-PCR primer design is illustrated in Fig. 3 and Table S4. The primer pairs of pJ277-pJ154, pJ277-pJ281, pJ283-pJ160, and pJ283-pJ281 were used to specifically detect unedited *Slc35f2* endogenous mRNA expression, edited *Slc35f2* endogenous mRNA expression, unedited *Slc35f6* endogenous mRNA expression and edited *Slc35f6* endogenous mRNA expression, respectively. Sequences of the forward primers correspond to the upstream exon not included in the dRT while sequences of the reverse primers correspond to the 3’ region of the last exons for unedited *Slc35f2* and *Slc35f6* alleles and to the APEX2 sequence of exogenous expression cassette for edited *Slc35f2* and *Slc35f6* alleles, respectively. The primer pair of dn736-dn737 was used for RT-PCR to amplify a 123-bp fragment from endogenously expressed *Gapdh* transcript as a positive control. RT-PCR was performed with 2 μl of the reverse transcription positive reactions and Phusion High Fidelity DNA polymerase (NEB, Ipswich, MA) following manufacturer’s protocols on an Applied Biosystems 2720 Thermal Cycler. Control RT-PCR reactions were performed with 2 µl of the reverse transcription negative reactions, containing only total RNA without cDNA synthesis, to detect any false positive result due to potential genomic DNA contamination. Control RT-PCR reactions resulted in no product amplification in all samples (data not shown) indicating no genomic DNA contamination in samples.

Aliquots of individual RT-PCR reactions with one pair of primers were analyzed on agarose gels, and then aliquots of equal volumes of relevant RT-PCR reactions were combined and loaded in a single gel to create a composite image (Fig. 3). Products of RT-PCR with expected size were amplified from cDNAs generated from each tissue sample of edited and wildtype animals, and then sequenced by Sanger Sequencing provided by Genewiz (genewiz.com, South Plainfield, NJ) to confirm correct splice junctions between adjacent exons and accurate transition from endogenous coding sequence to the in-frame inserted exogenous expression cassette. Sequence data were analyzed by Geneious Prime software (geneious.com, Auckland, New Zealand).

### SURVEYOR Assay for iCAP edited MED13L allele

In order to evaluate the efficiency of the mutation correction by exchange of exon 20 mediated by iCAP genome editing, the SURVEYOR nuclease assay was performed to estimate the reduction of mutant alleles in edited cell populations in comparison with those in un-edited patient cells. Genomic DNAs were extracted from WI-38 fibroblasts [*MED13*L^exon20(+/+)^], the MED13L Syndrome patient cells [*MED13L*^exon20(+/−)^], 4-1-2 cells [*MED13L*^exon20(+/−, +/+)^], and 5-1-2 cells [*MED13L*^exon20(+/−, +/+)^]. Nested PCRs were performed individually with genomic DNAs from each group of cells. An approximate 593 bp PCR fragment was generated using a forward primer F1 (pJ414: TATCAGATGTGTAGGCTTGGGCAG) corresponding to the intron19-20 sequence immediately 5’ of exon20 and a reverse primer R1 (pJ355: GTCTCCTTTCAGACTGATTCCATG) with the corresponding intron20-21 sequence downstream of the region presented in the dRT, therefore excluding amplification from randomly integrated dRTs. The resultant PCR fragment would contain the exon20 sequences with and without the single thymine duplication as well as the intron20-21 sequence with the 3’ cleavage sites (for cognate Cas9 or Cas12a cleavage) (Fig. 6a). To exclude any mismatch occurring at the 3’ cleavage sites from the assay, the 593 bp PCR products were used as templates for a second round of PCR to produce a 270 bp fragment with the same forward primer F1 (pJ414) and a reverse primer R2 (pJ415: AAAAGAAGGGATGGGACGGATTTG) corresponding to the intron sequence 3’ of exon20 but distantly 5’ to the 3’ cleavage sites. This smaller PCR fragment would only contain the single nucleotide duplication in exon20 as a mismatch location (Fig. 6a) and thus used for the SURVEYOR assay.

In Brief, the 270-bp PCR products from each group of cells were column purified and then an aliquot of 0.5 μg was heated at 100°C for 10 minutes for denaturing, followed by a slow cool for 1hour at room temperature to allow re-hybridization to form WT homoduplex, mutant homoduplex and WT/mutant heteroduplex fragments. The re-hybridized PCR products were used for the SURVEYOR assay according to manufacturer’s protocols (IDT, Coralville, IA), and results were analyzed on a 2% agarose gel (Fig. 6b). Cleavage by SURVEYOR nuclease at the mismatch site generates smaller fragments predicted as 156-bp and 114-bp bands. To quantify the ratio of mutant and non-mutant allele, the bands of 270 bp (*a*), 156 bp (*b*) and 114 bp (*c*) fragments in an agarose gel were visualized and the intensities of each band were measured on a Gel Logic 112 Digital Imaging System with Molecular Imaging Software 5.0 (mi.carestreamhealth.com, Rochester, NY). Values for the relative intensities of individual gel bands were obtained (Table S2). The indel rate for each group of cell populations was determined by the formula, [1- (1- (*b*+*c*)/(*a*+*b*+*c*))^1/2^] x100 (Ran et al, 2013), where *a* is the intensity of un-cleaved PCR product, and *b* and *c* are the intensities of cleavage products in each group of cell populations. A reduced indel rate in a group of iCAP edited patient cell populations reflects reductions of the mismatch as a result of the elimination of a thymine duplication from *MED13L* mutant alleles. By a cross reference to the indel rate in un-edited patient cells, mismatch reduction percentage was estimated by the formula, [1- (*e/m*)] x100, where *e* is an indel rate in non-clonal edited patient cell populations and *m* is the indel rate in un-edited *MED13L^S1497F^* patient cells, an indirect reflection of iCAP editing efficiency.

### Detection of cellular oxidative stress with dihydroethidium (DHE) probe

2 x 10^4^ cells collected from each cell group of WI-38, unedited and edited *MED13L^S1497F^* patient fibroblasts were plated in wells of 12-well plates respectively and cultured in the same condition as described above for 24 hours. Cells were then harvested by trypsinization followed by centrifugation, resuspended in 300 µl of PBS containing 10 µM DHE (Thermo Fisher Scientific, Waltham, MA) and maintained in the dark at 37°C for 30 min. Fluorescence was measured by BD Accuri^TM^ C6 flow cytometer. All measurements were conducted for three independent cultures of each cell group and with 10,000 events. Data were analyzed by FlowJo (Tree Star, Inc., Ashland, OR) and statistical analysis was performed using Student’s t test with p ≤ 0.05 considered significant.

### Off-target effect analysis

The top 4 or 5 off-target (OT) sites for each gRNAs selected for the study were chosen based on predication by the online gRNA design tool Benchling (www.benchling.com). The “position” of the OT sites designates their predicted cut-site at genomic location proceeded by a chromosome number reported by the Benchling software (Table S3). PCR primers were designed to amplify an approximately 600 bp product with 300 bp 5’ and 3’ flanking the predicted cut-site and listed in Table S4. PCR amplicons produced from experimental (edited) and control (unedited or wildtype) genomic DNA samples were Sanger Sequenced by GeneWiz sequencing service. DNA sequences were then compared & analyzed by using the online <u>I</u>nference of <u>C</u>RISPR <u>E</u>dits (ICE) algorithm tool (Conant, et al, 2022) (https://www.synthego.com/products/bioinformatics/crispr-analysis). A report of ICE analysis provides an R^2^ value and indel rate. The R^2^ value between 0-1.0 indicates “goodness of fit” for the alignment of the sequence data and model created for each comparison between experimental and control samples as well as for a value that predictes indel percentage or frequency of indels in the experimental sample compared to control. An acceptable R^2^ value is >0.8, and an indel percentage of >5% indicates a true indel.

## ACKNOWLEDGEMENTS

We thank Jeremy Wang, Nicolae (Adrian) Leu and Stephanie Sterling at University of Pennsylvania School of Veterinary Medicine for providing mouse production services.

## AUTHOR CONTRIBUTIONS

P.J., Y.L., and R.S. conceived the projects to modify the mouse and human genes. P.J., K.K and S.C. conceived the original idea of iCAP. P.J., K.K., K.Z., C.Q. and L.G. performed experiments. P.J. and K.K. designed gene editing schemes. K.K. contributed to CRISPR/Cas molecular component production, molecular cloning and genotyping. P.V.H. identified the patient carrying *MED13L^S1497F^* mutation and established the cell line of *MED13L^S1497F^* fibroblasts. P.J., Y.L., P.V.H. and R.S. contributed to study design and collaboration setup. P.J. drafted the manuscript. R.S., K.K., S.C. and P.J. contributed to manuscript revision.

## SUPPLEMENTARY FIGURES

**Figure S1.**
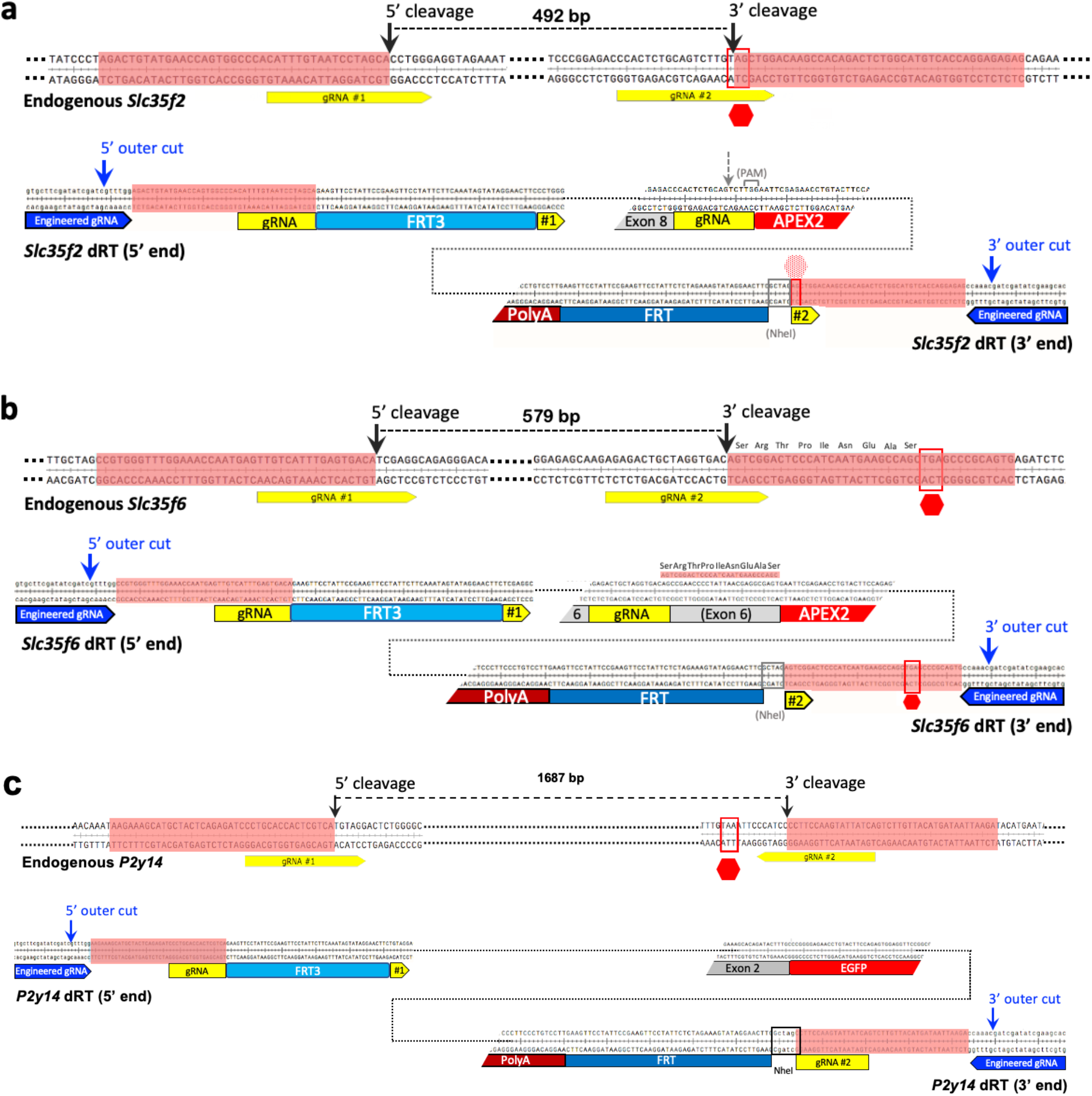
DNA sequences at up- (5’) and downstream (3’) cleavage sites in loci of mouse genome and in edited isogenic fragment on dRT. (**a**) DNA sequence at 5’ and 3’ cleavage sites in *Slc35f2* gene locus (top). Two gRNA target sequences (indicated by arrowed yellow bars) flank a 492 bp genomic target segment (GTS) containing partial intron 7-8 and coding sequence of exon 8. Cas9 cleavages (black arrows) excise GTS and expose end sequences of 40-bp as overhang substrate sequence (OSS) shaded in salmon. Edited sequence structures in EIF on *Slc35f2* dRT (bottom). Two genomic gRNA target sequences are disrupted by insertions of a FRT3 site and an APEX2-Cre expression cassette, respectively. Cas9 cleavages (blue arrows) at engineered gRNA target sites excise the 4386-bp EIF, which contains GTS with precisely inserted exogenous sequences, and expose the EIF’s end sequences of 40-bp OSS, homology to those at the corresponding genomic ends. A PAM site (in parentheses) is unintentionally created in the truncated 3’ gRNA target sequence at the junction of exon8-APEX2. (**b**) DNA sequence at 5’ and 3’ cleavage sites in *Slc35f6* gene locus (top). Two gRNA target sequences (indicated by arrowed yellow bars) flank a 579 bp GTS containing partial intron 5-6 and coding sequence of exon 6. Edited sequence structures in EIF on *Slc35f6* dRT (bottom). Two genomic gRNA target sequences are disrupted by insertions of a FRT3 site and APEX2-Cre expression cassette, respectively. As the 3’ gRNA target sequence is coding DNA for last 9 amino acids, non-sense mutations are made to the sequence and PAM site at the junction of exon6-APEX2 to avoid cleavage. Cas9 cleavages (blue arrows) at engineered gRNA target sites excise the 4501-bp EIF. (**c**) DNA sequence at 5’ and 3’ cleavage sites in *P2y14* gene locus (top) and edited sequence structures in EIF on *P2y14* dRT (bottom). Cas9 cleavages (blue arrows) at engineered gRNA target sites excise the 5482-bp EIF. Bases in salmon, OSS; bases in red frame, stop codon; faded stop sign, location of original stop codon; vertical arrows, Cas9 cleavage sites.

**Figure S2.**
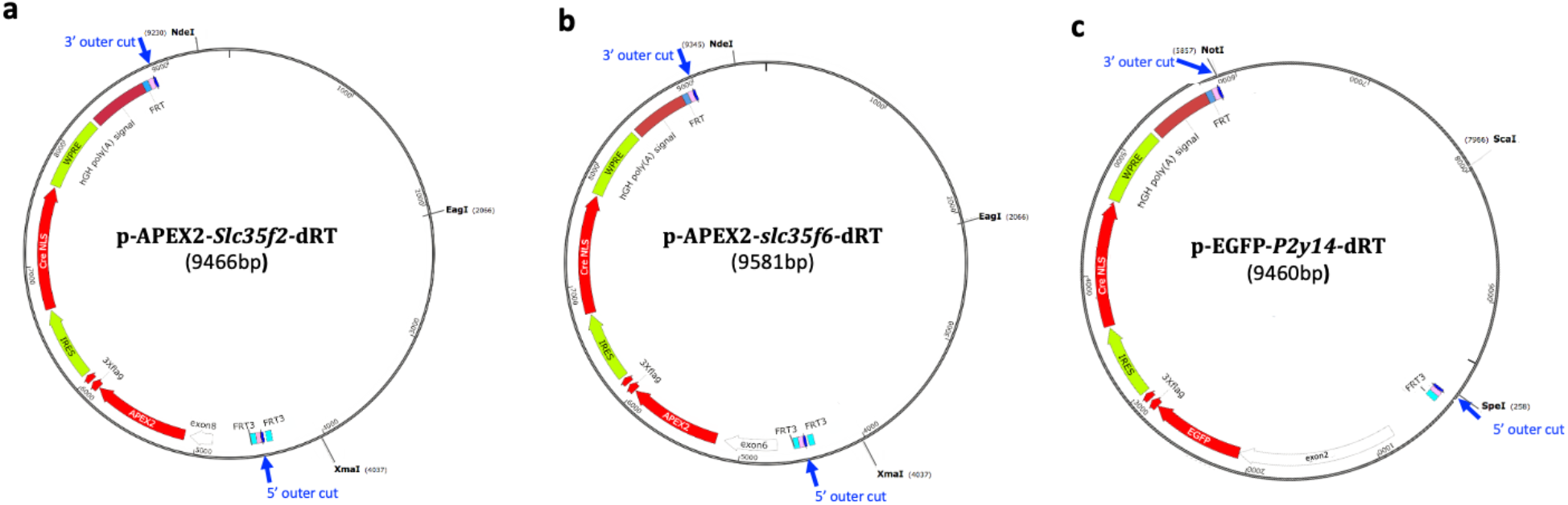
Maps of plasmids containing dRT fragments for mouse gene editing. **(a)** *Slc35f2* dRT plasmid. (**b)** *Slc35f6* dRT plasmid. XmaI-NdeI-EagI triple restriction enzyme digestions produce Xmal-NdeI dRT fragments (5193-bp *Slc35f2* dRT and 5308-bp *Slc35f6* dRT) and two smaller EagI-XmaI and EagI-NdeI vector backbone fragments for clear separation of dRT fragment from vector backbone. (**c)** *P2y14* dRT plasmid. SpeI-NotI-ScaI triple restriction enzyme digestions produce SpeI-NotI dRT fragment (5599-bp) and two smaller ScaI-SpeI and ScaI-NotI vector backbone fragments.

**Figure S3.**
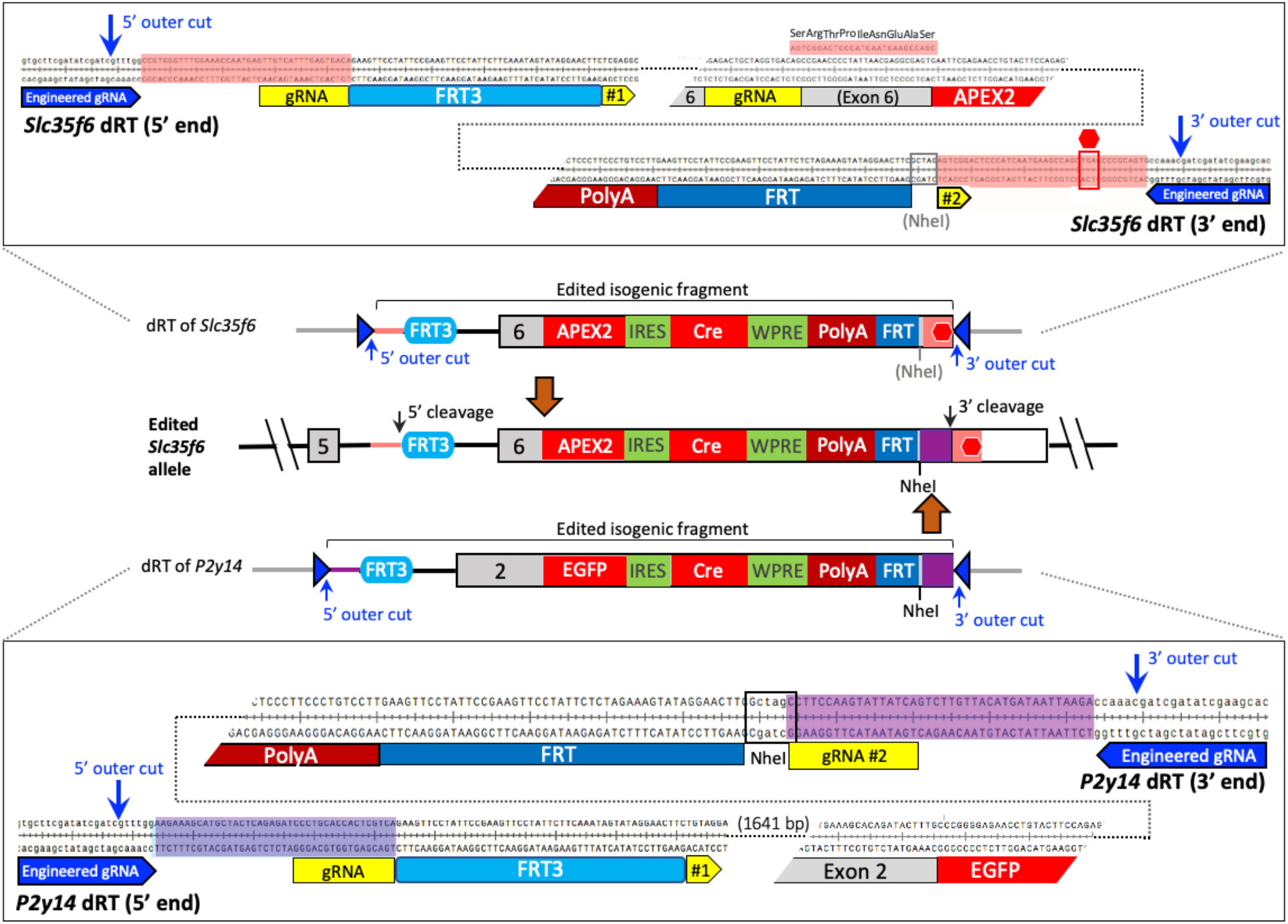
Exchange between 3’ ends of two different edited isogenic fragments by gene conversion. Schematic and sequence of *Slc35f6* dRT (top panel). The 5’ portion of EIF from *Slc35f6* dRT contributed to the corresponding portion of pasted EIF in the edited *Slc35f6* allele (middle). Schematic and sequence of *P2y14* dRT (bottom panel). The 3’ portion of EIF from *P2y14* dRT contributed to the corresponding portion of pasted EIF in the edited *Slc35f6* allele as both *P2y14* dRT and *Slc35f6* dRT were co-introduced into zygote. The exchange between the identical 3’ portions of the two different EIFs might be an outcome of nonallelic gene conversion. Sequence in salmon, OSS flanking *Slc35f6* EIF; sequence in purple, OSS flanking *P2y14* EIF; NheI in grey, the site lost; NheI in black, the site retained.

**Figure S4.**
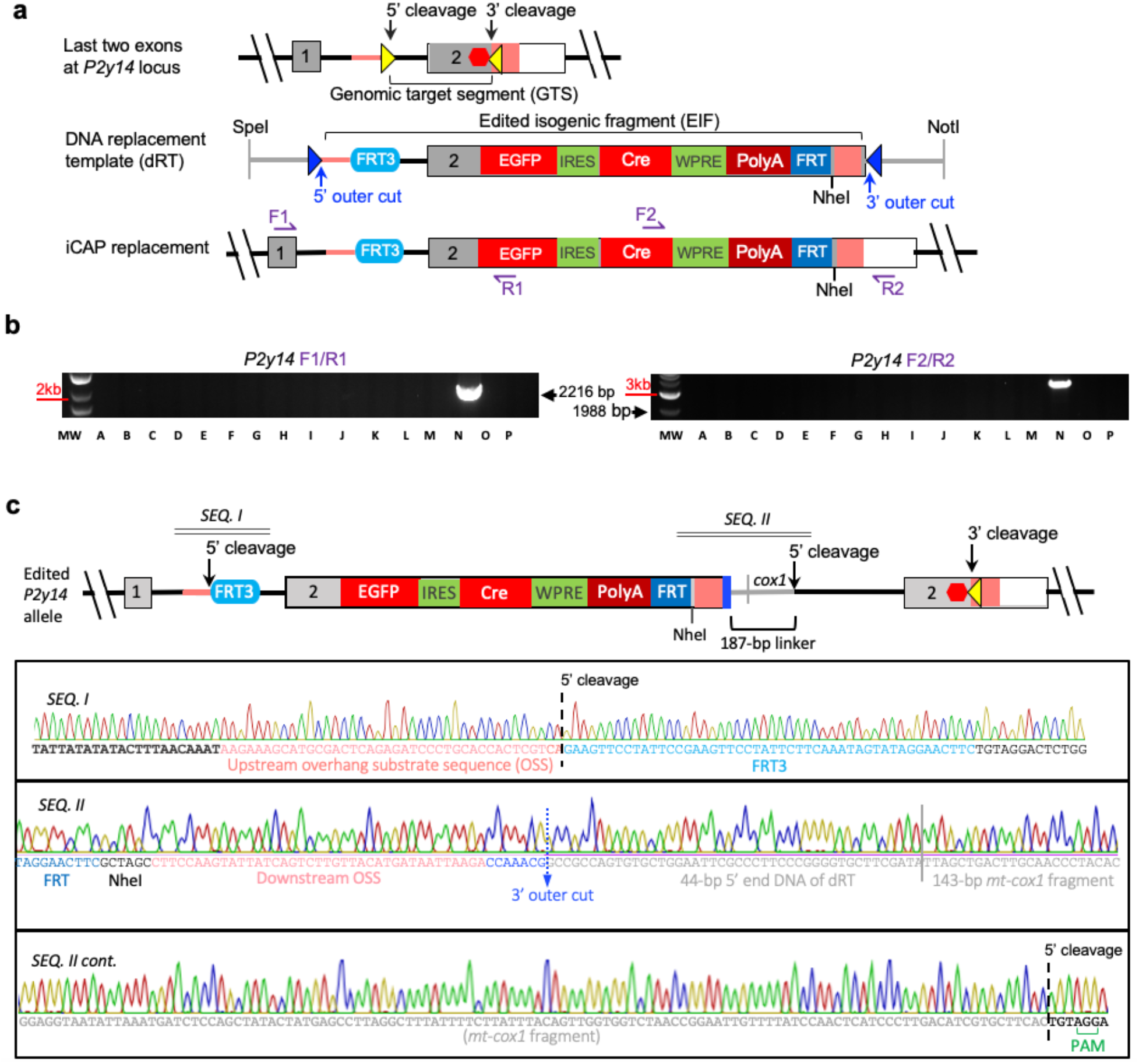
Edited *P2y14* allele in Animal N. **(a)** Depicted strategy of iCAP genome editing mediated by CRISPR/Cas9. (**b**) Genotyping results of 16 mice. An expected PCR product of 2,216-bp spanning the junction of 5’ paste site and a larger than predicted (1,988-bp) PCR fragment spanning the junction of 3’ paste site indicates that Animal N is a candidate containing iCAP edited allele. (**c**) Schematic showing targeted integration of an edited isogenic fragment (EIF) excised from *P2y14* dRT at the 5’ cleavage site (top) and DNA sequence of edited *P2y14* region (sequence panel). The end-joining between the 5’ genomic DSB end at 5’ cleavage site and the 5’ end of *P2y14* EIF is error free (*SEQ. I)*. The 3’ end of *P2y14* EIF was properly exposed by Cas9 cleavage but the 3’ genomic end is in fact the PAM proximal DSB end at the same 5’ cleavage site rather than at the 3’ cleavage site where no cleavage occurs. These two relatively undefined ends are bridged by a linker of 187 bp sequence comprised of a 44 bp vector backbone section that is the sequence at the 5’ end of dRT to mask OSS and a 143 bp sequence identical to part of mitochondrial cytochrome oxidase subunit I (*Cox1*) gene (*SEQ. II)*. Black bold bases, sequences not included in dRT; NheI, restriction site.

**Figure S5.**
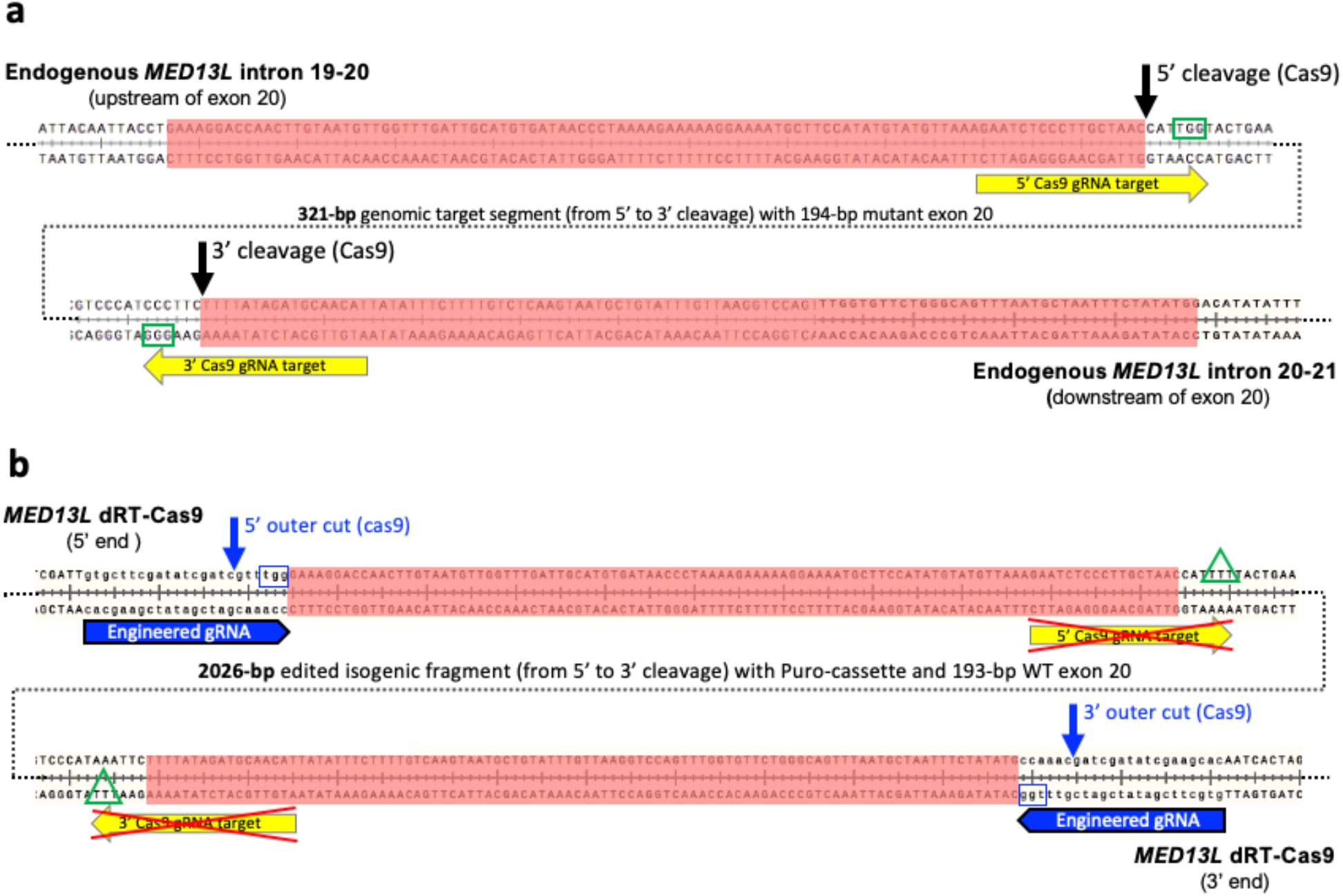
Sequence structures at 5’ and 3’ cleavage sites in *MED13L* locus and at engineered gRNA target sites in iCAP Cas9 dRT. **(a)** Sequence structures at 5’ and 3’ cleavage sites in *MED13L* locus. These two cleavage sites flanking exon 20 are used for excision of GTS. Bases highlighted in salmon, 5’ 100-bp and 3’ 100-bp OSS respectively; yellow-boxed arrow, gRNA target sequences with PAM site in green frame; black arrows, locations of DSB induced by Cas9; black dotted line, the intervening sequence (GTS) between DSBs. **(b)** Sequence structures at engineered gRNA target sites in iCAP Cas9 dRT. The engineerd gRNA target sites are used for excision of the edited isogenic fragment (EIF) from dRT. Blue-boxed arrow, engineered gRNA target sequences with PAM site in blue frame; vertical blue arrow, locations of DSB induced by Cas9; bases highlighted in salmon, 5’ 100-bp and 3’ 100-bp OSS respectively; crossed yellow-boxed arrow, disabled gRNA target sequences with mutated PAM site in green delta frame; black dotted line, the EIF between two outer cuts.

**Figure S6.**
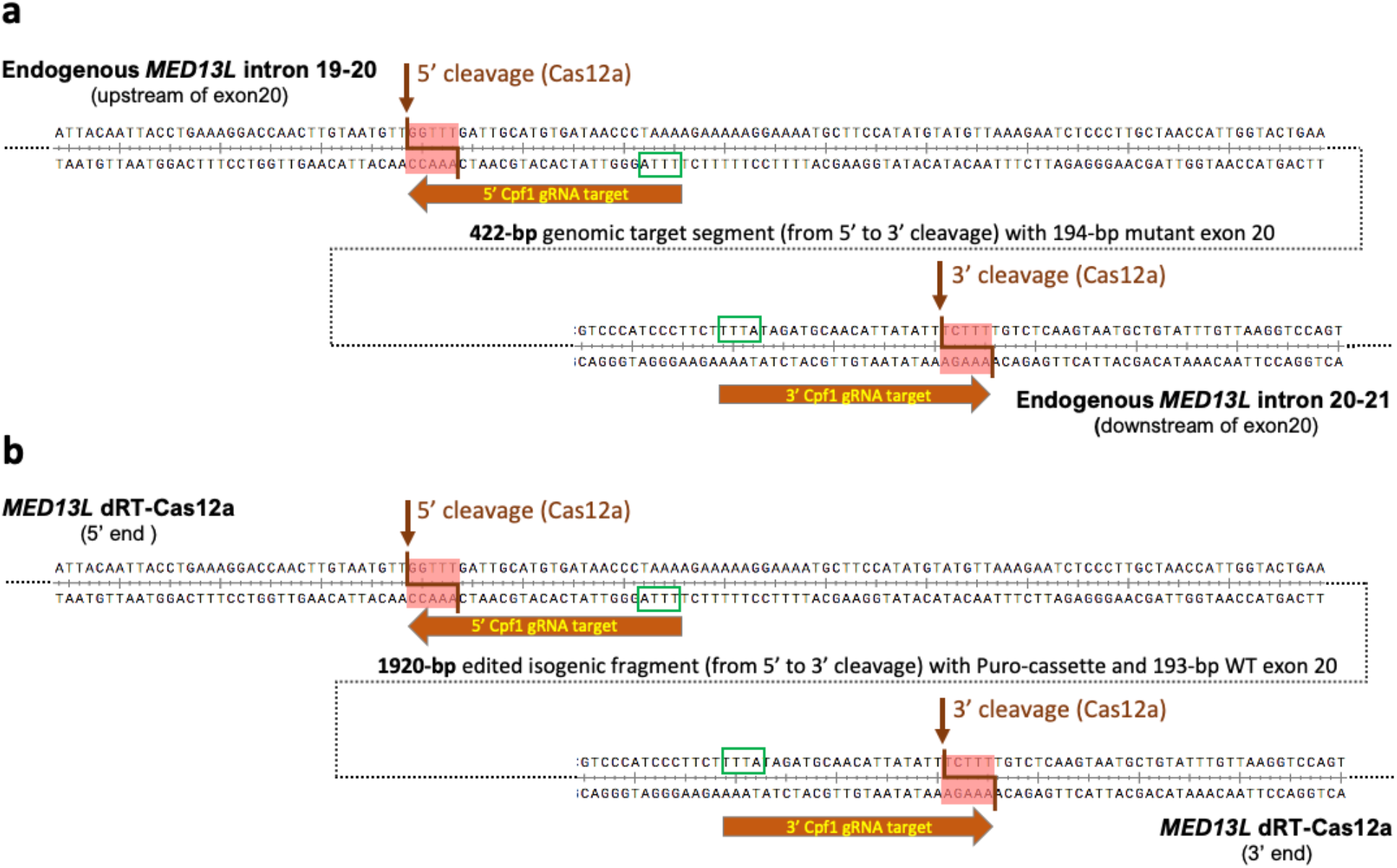
Sequence structures at 5’ and 3’ cleavage sites in *MED13L* locus and in iCAP Cas12a dRT. **(a)** Sequence structures at 5’ and 3’ cleavage sites in *MED13L* locus. These two cleavage sites flanking exon 20 are used for excision of GTS. Dark orange boxed arrow, gRNA target sequences with PAM site in green frame; brown arrow and elbow line, locations of staggered DSB induced by Cas12a; black dotted line, the intervening sequence (GTS) between DSBs. **(b)** Sequence structures at the 5’ and 3’ cleavage sites that are fully retained in iCAP Cas12a dRT. The same 5’ and 3’ cleavage sites are used for excision of the edited isogenic fragment (EIF) from dRT. Dark orange boxed arrow, gRNA target sequences with PAM site in green frame; brown arrow and elbow line, locations of staggered DSB induced by Cas12a; black dotted line, the EIF between DSBs.

**Figure S7.**
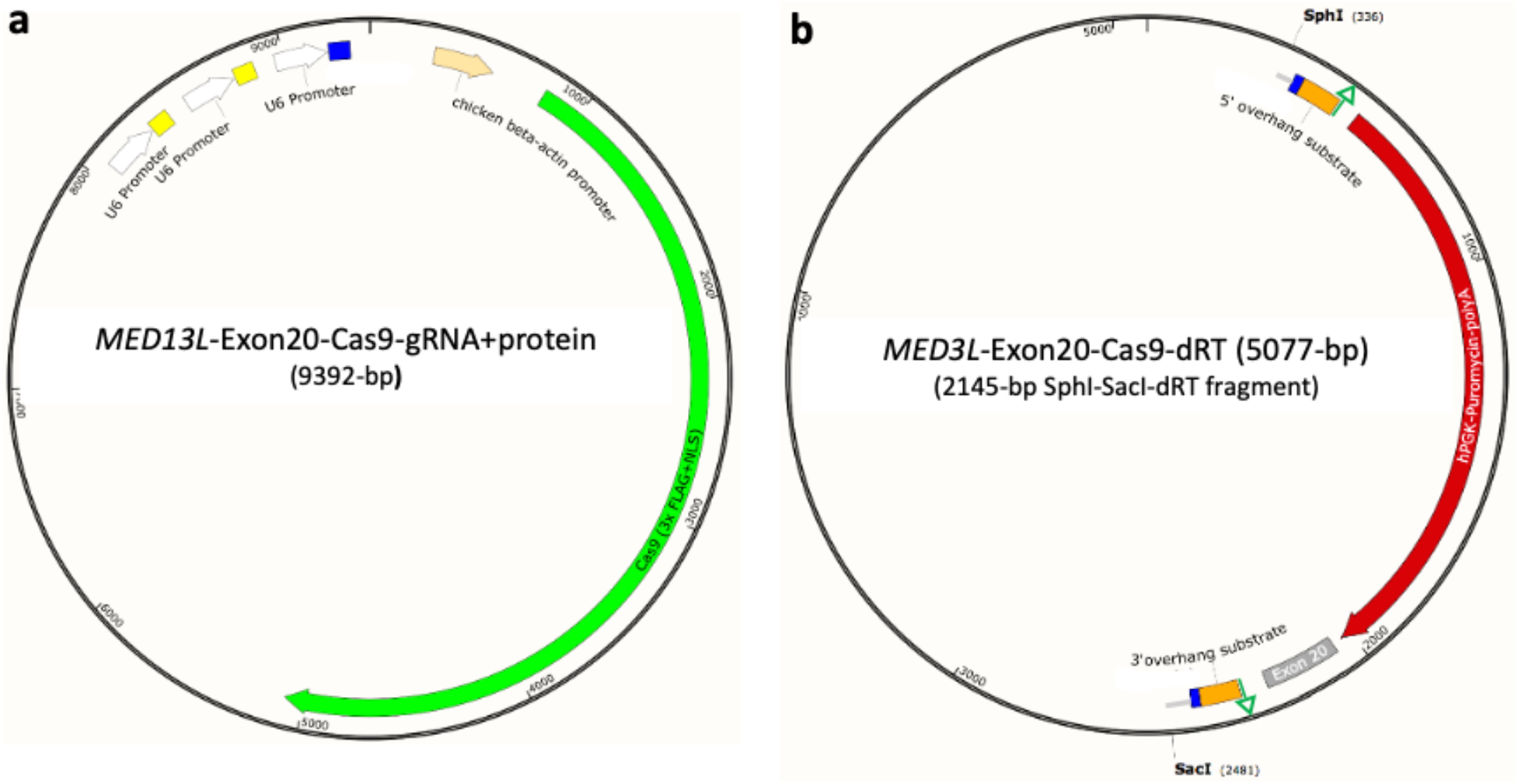
Map of CRISPR/Cas9 expression plasmid and dRT-containing plasmid for *MED13L* editing. **(a)** Plasmid for expression of CRISPR/Cas9. In addition to Cas9 gene, the plasmid contains three crRNA genes (yellow and blue boxes) for expressions of three different sgRNAs. Two of the sgRNAs target the 5’ and 3’ gRNA sites flanking exon 20 of the *MED13L* gene for inducing 5’ and 3’ cleavages to excise GTS, and the third one targets the engineered gRNA target site for inducing “outer cuts” to excise EIF from the dRT. **(b)** Plasmid containing dRT used for Cas9 mediated iCAP editing. SphI-SacI restriction digestion produces the 2145-bp dRT fragment used for transfection. Blue boxes, engineered gRNA target sites; green arrows, mutated PAM sites associated with genomic gRNA target sites; grey bar, vector backbone sequence.

**Figure S8.**
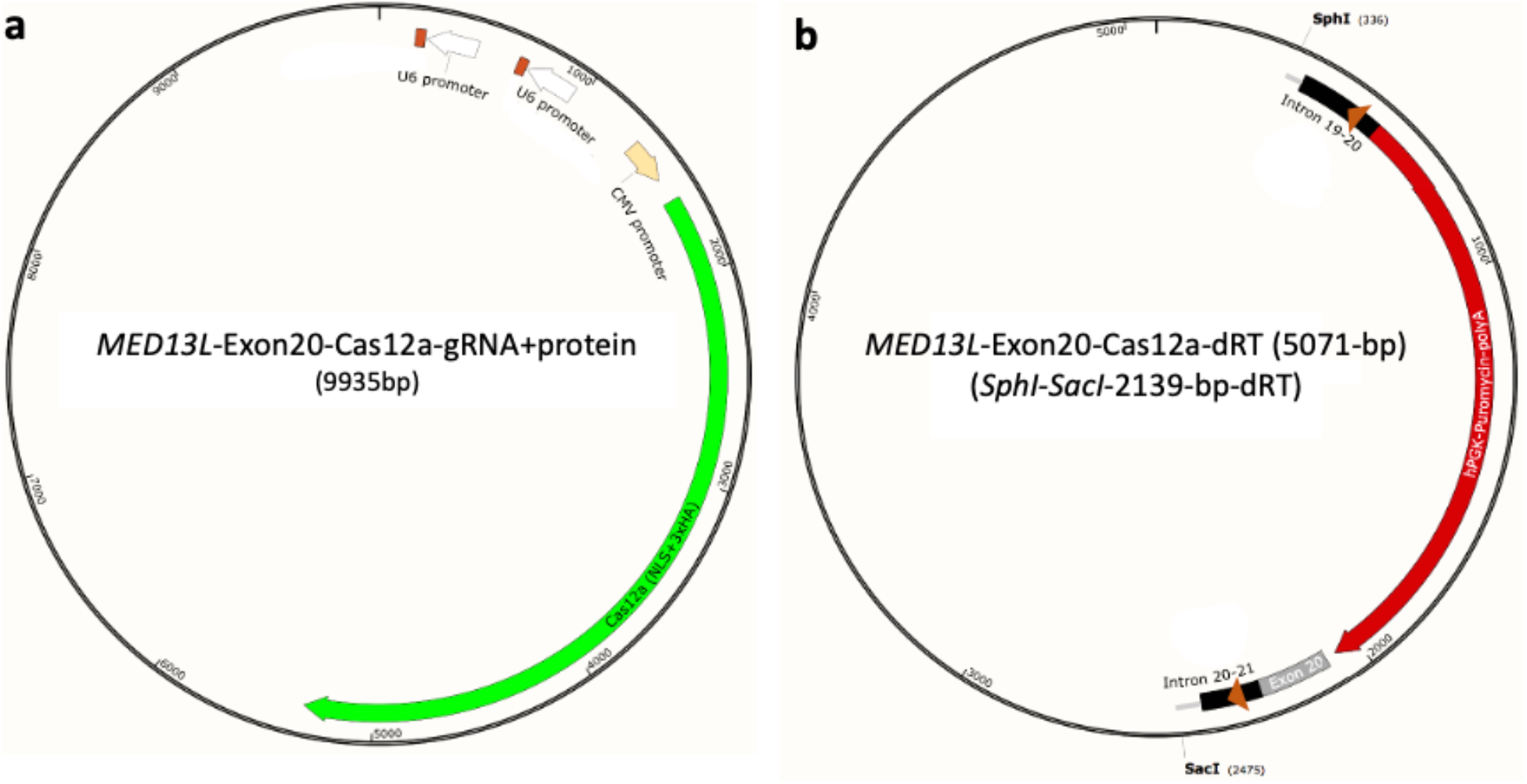
Map of CRISPR/Cas12a expression plasmid and dRT-containing plasmid for *MED13L* editing. **(a)** Plasmid for expression of CRISPR/Cas12a. In addition to Cas12a gene, the plasmid contains two crRNA genes (orange box). The expressed sgRNAs target the 5’ and 3’ gRNA sites flanking exon 20 for inducing 5’ and 3’ cleavages to excise both GTS from the endogenous *MED13L* gene and EIF from the dRT. (**b)** Plasmid containing dRT used for Cas12a mediated iCAP editing. SphI-SacI restriction digestion produces the 2139-bp dRT fragment used for transfection. Orange arrowheads, genomic 5’ and 3’ gRNA target sites; grey bar, vector backbone sequence.

## SUPPLEMENTARY TABLES

**Table S1.**
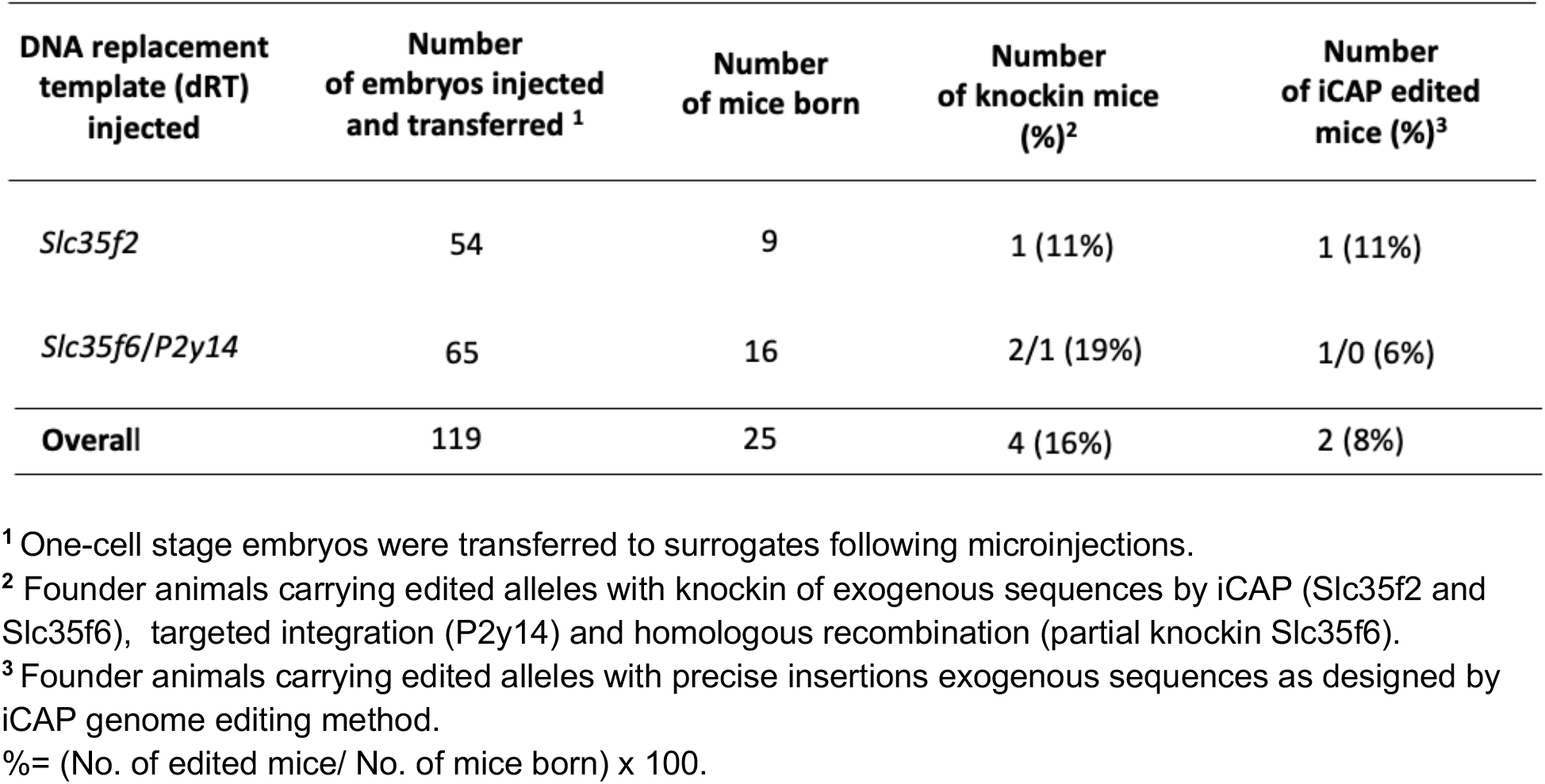
Production of iCAP edited animals.

| DNA replacement template (dRT) injected | Number of embryos injected and transferred <sup>1</sup> | Number of mice born | Number of knockin mice (%)<sup>2</sup> | Number of iCAP edited mice (%)<sup>3</sup> |
| --- | --- | --- | --- | --- |
| *Slc35f2* | 54 | 9 | 1 (11%) | 1 (11%) |
| *Slc35f6/P2y14* | 65 | 16 | 2/1 (19%) | 1/0 (6%) |
| **Overall** | **119** | **25** | **4 (16%)** | **2 (8%)** |

<sup>1</sup> One-cell stage embryos were transferred to surrogates following microinjections.
<sup>2</sup> Founder animals carrying edited alleles with knockin of exogenous sequences by iCAP (*Slc35f2* and *Slc35f6*), targeted integration (*P2y14*) and homologous recombination (partial knockin *Slc35f6*).
<sup>3</sup> Founder animals carrying edited alleles with precise insertions exogenous sequences as designed by iCAP genome editing method.
%= (No. of edited mice/ No. of mice born) x 100.

**Table S2.** Measured values of relative band intensity in SURVEYOR assay.

|  | 4-1-2<br>(*Cas9*) | 5-1-2<br>(*Cas12a*) | Patient<br>cells | WT<br>Cells |
| --- | --- | --- | --- | --- |
| *a* | .5788 | .5944 | .4875 | 1 |
| *b* | .2766 | .2726 | .3332 | --- |
| *c* | .1445 | .1330 | .1793 | --- |

Note. 4-1-2 (*Cas9*), non-clonal edited *MED13L*<sup>S1497F</sup> patient cell populations by iCAP using Cas9; 5-1-2 (*Cas12a*), non-clonal edited *MED13L*<sup>S1497F</sup> patient cell populations by iCAP using Cas12a; Patient cells, *MED13L*<sup>S1497F</sup> patient fibroblasts; WT, WI-38 normal fibroblasts. Indel % = $[1 - (1 - (b+c)/(a+b+c))^{1/2}] \times 100$ , where *a* is the intensity of un-cleaved PCR product, and *b* and *c* are the intensities of cleavage products. Mismatch reduction percentage is estimated by the formula, $[1 - (e/m)] \times 100$ , where *e* is an indel rate in non-clonal edited patient cell populations and *m* is the indel rate in un-edited *MED13L*<sup>S1497F</sup> patient cells.

**Table S3.**
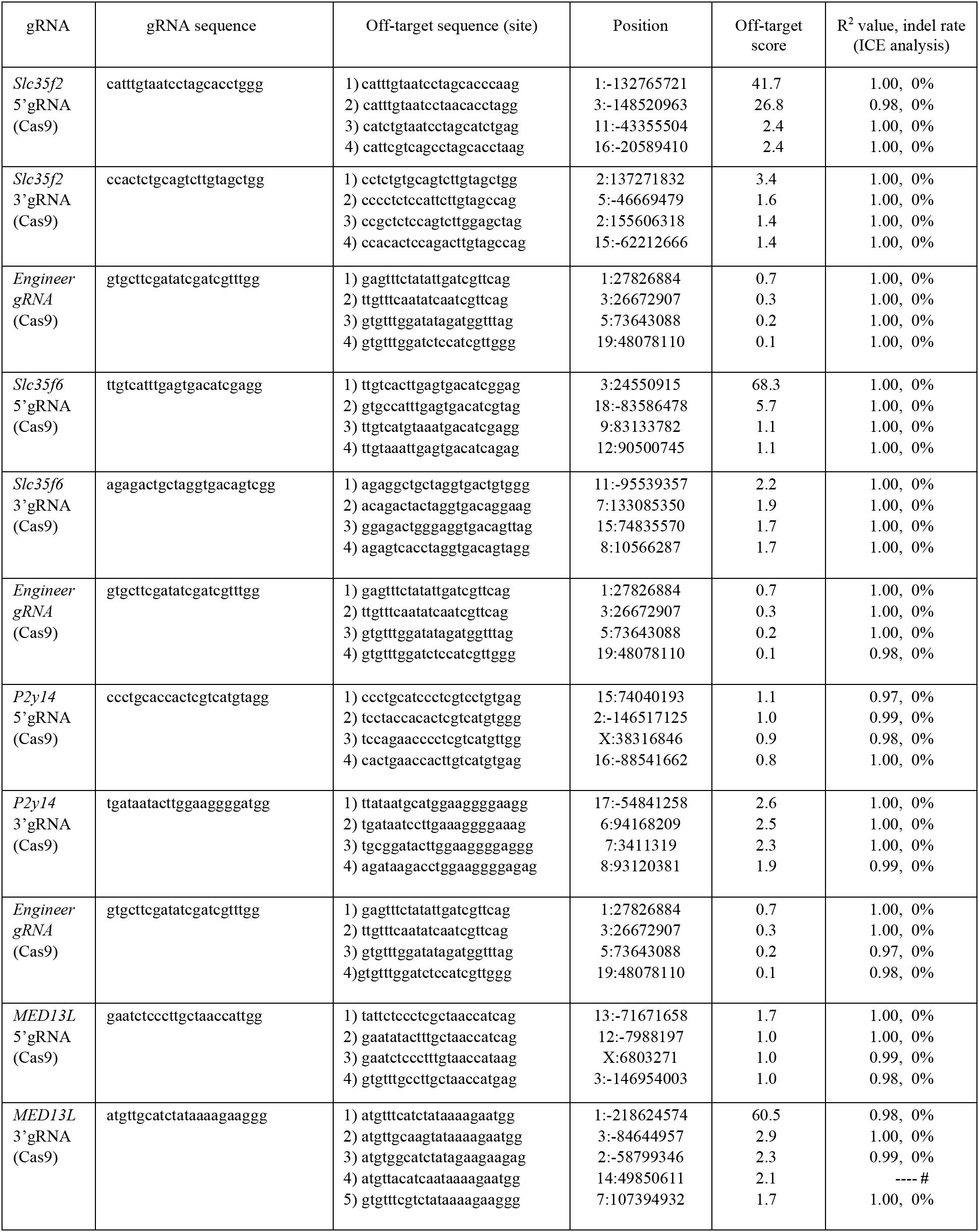
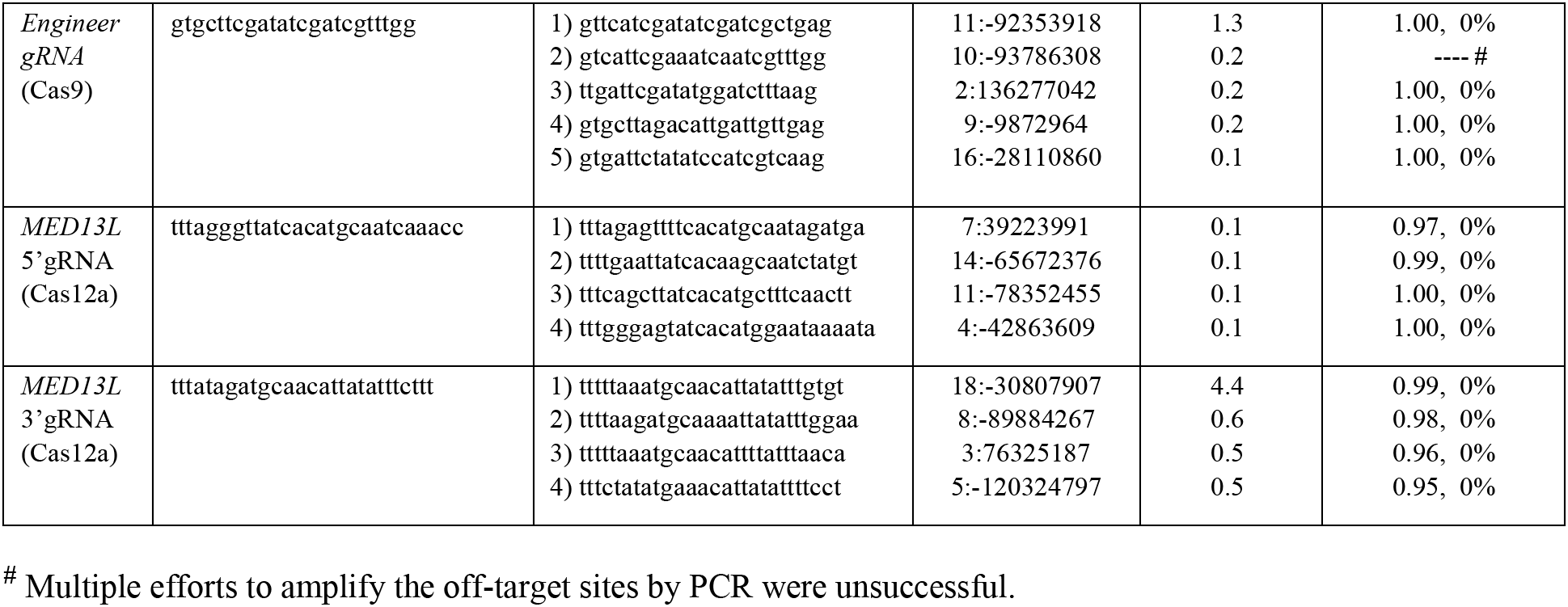
Analysis for Cas effector activities at top off-target sites.

**Table S4.**
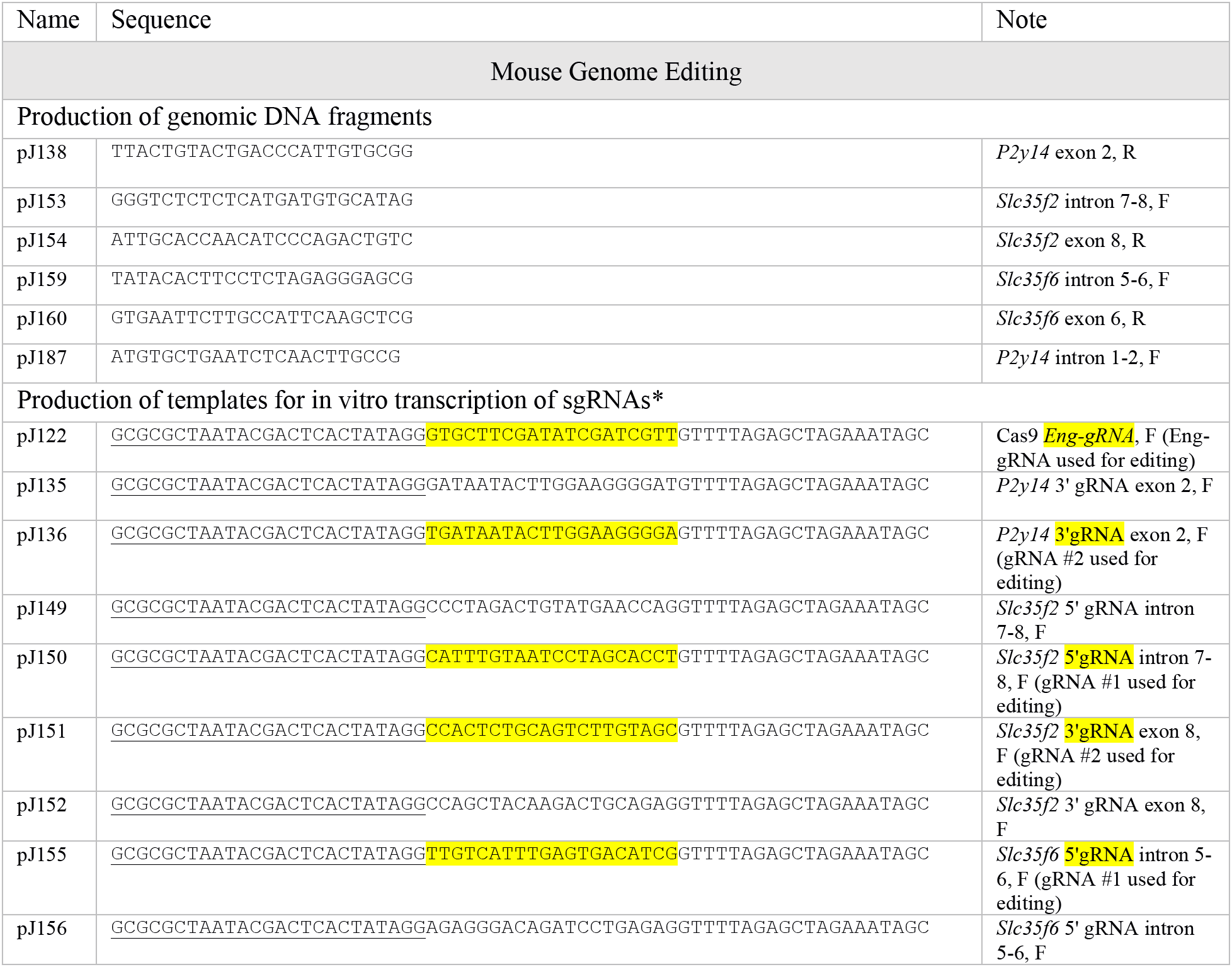
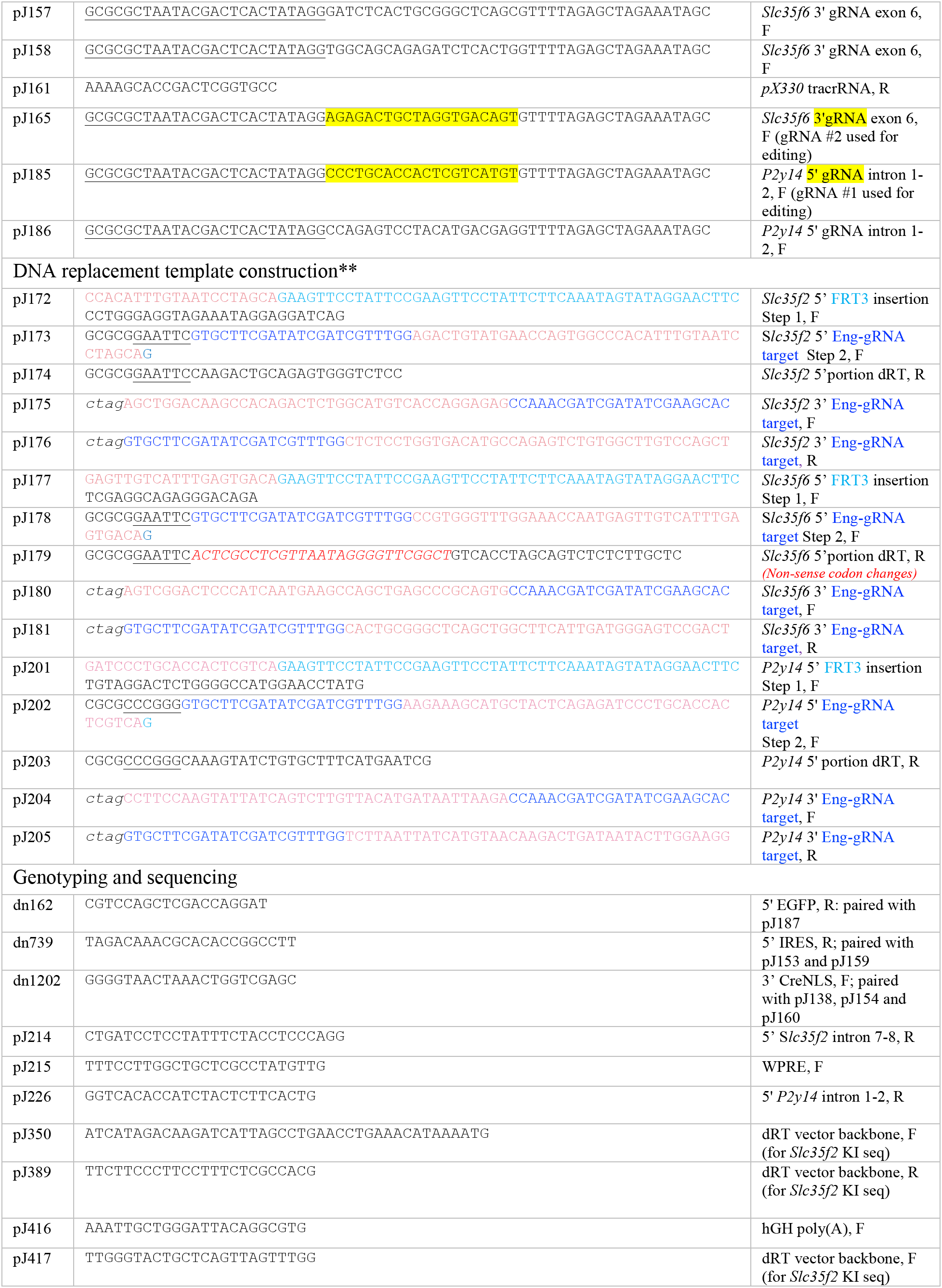
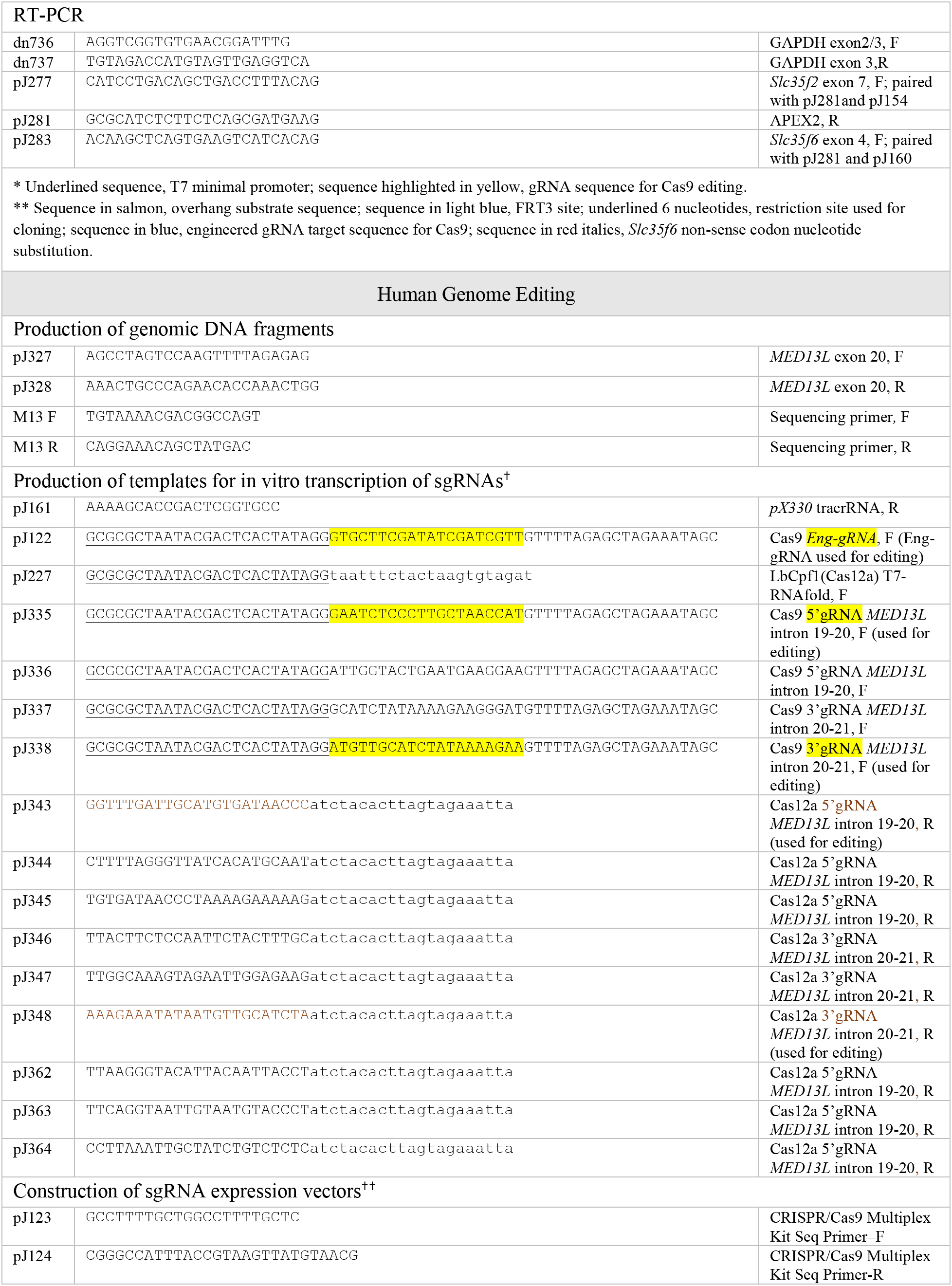
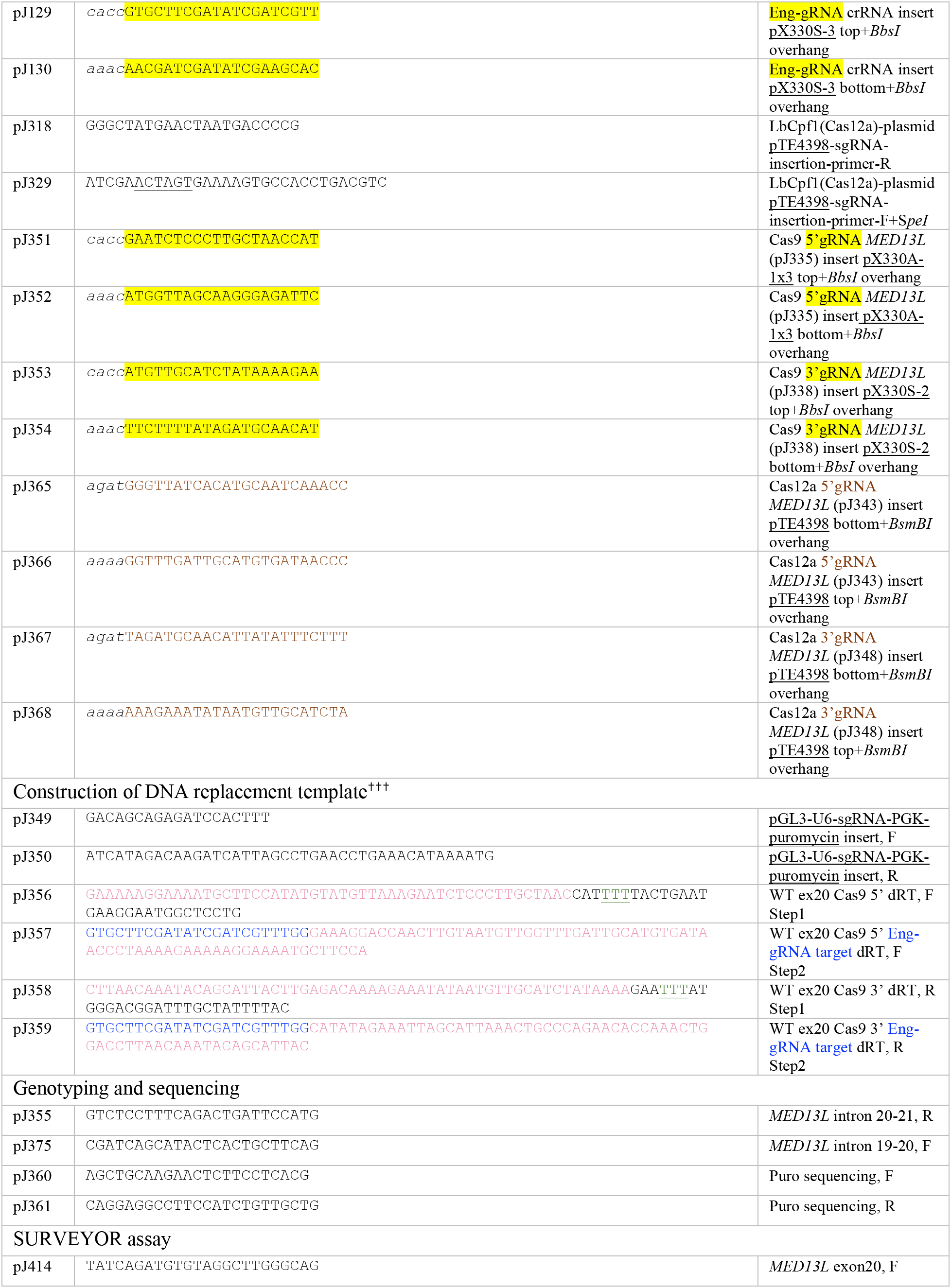
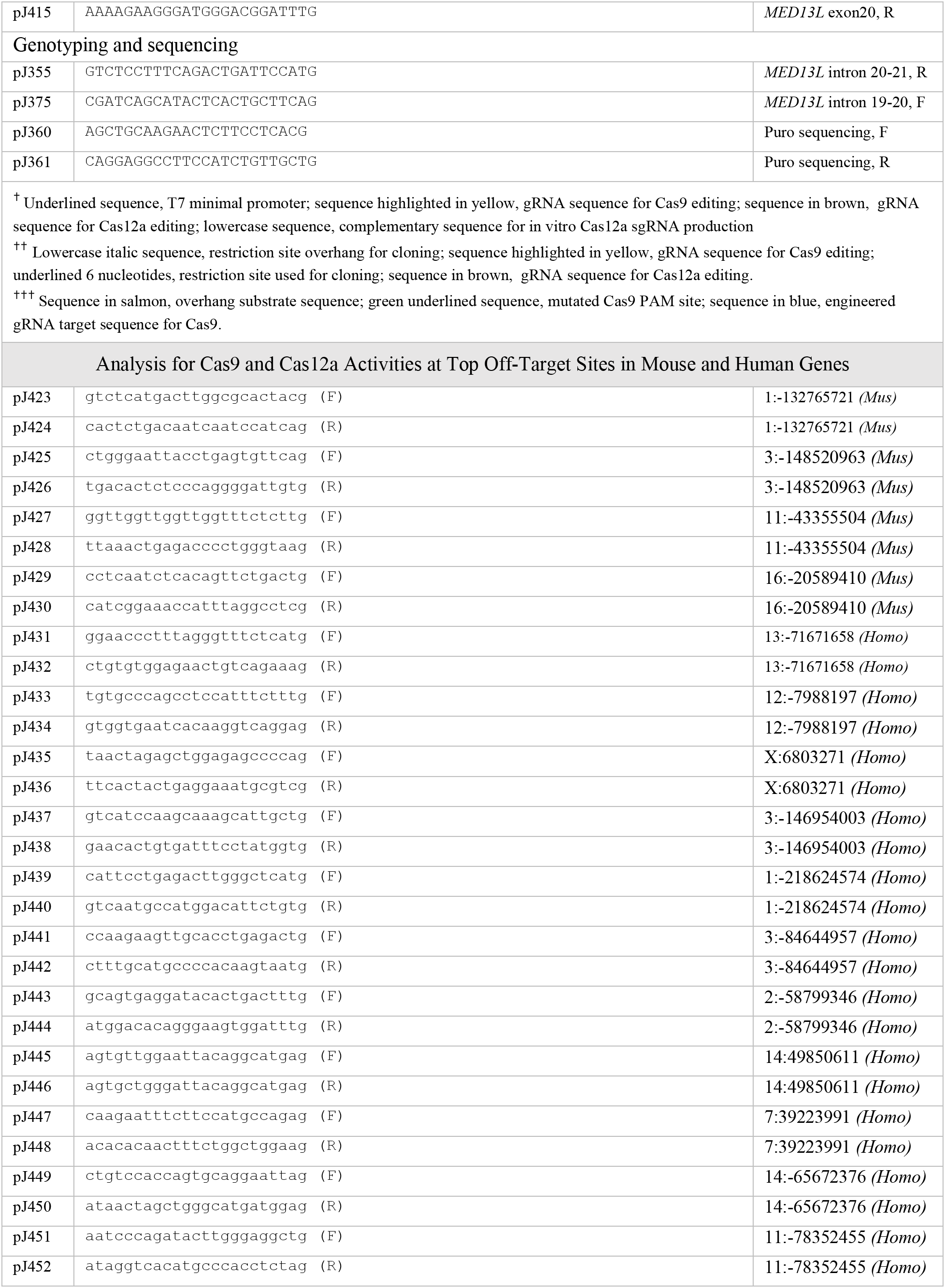
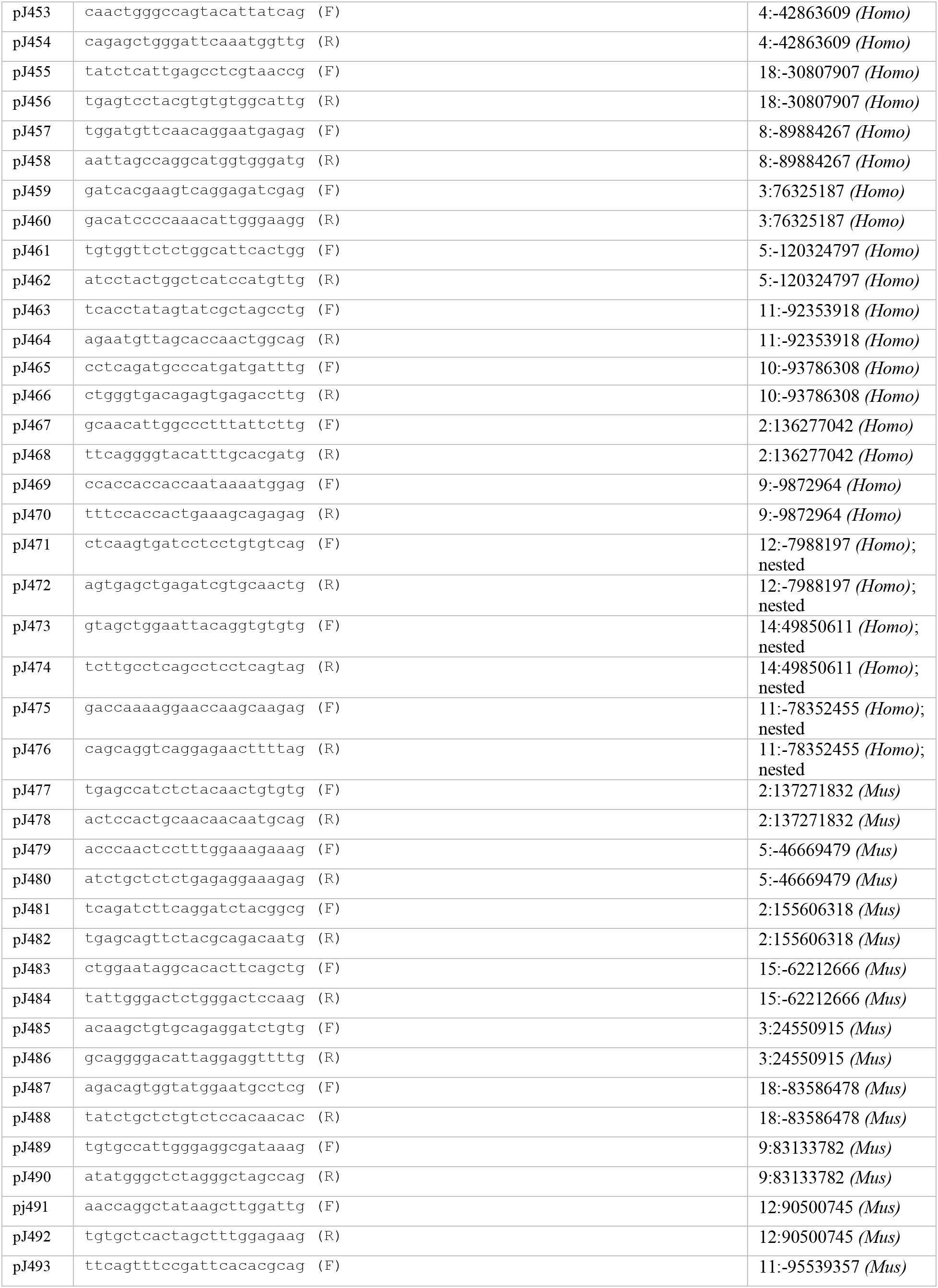
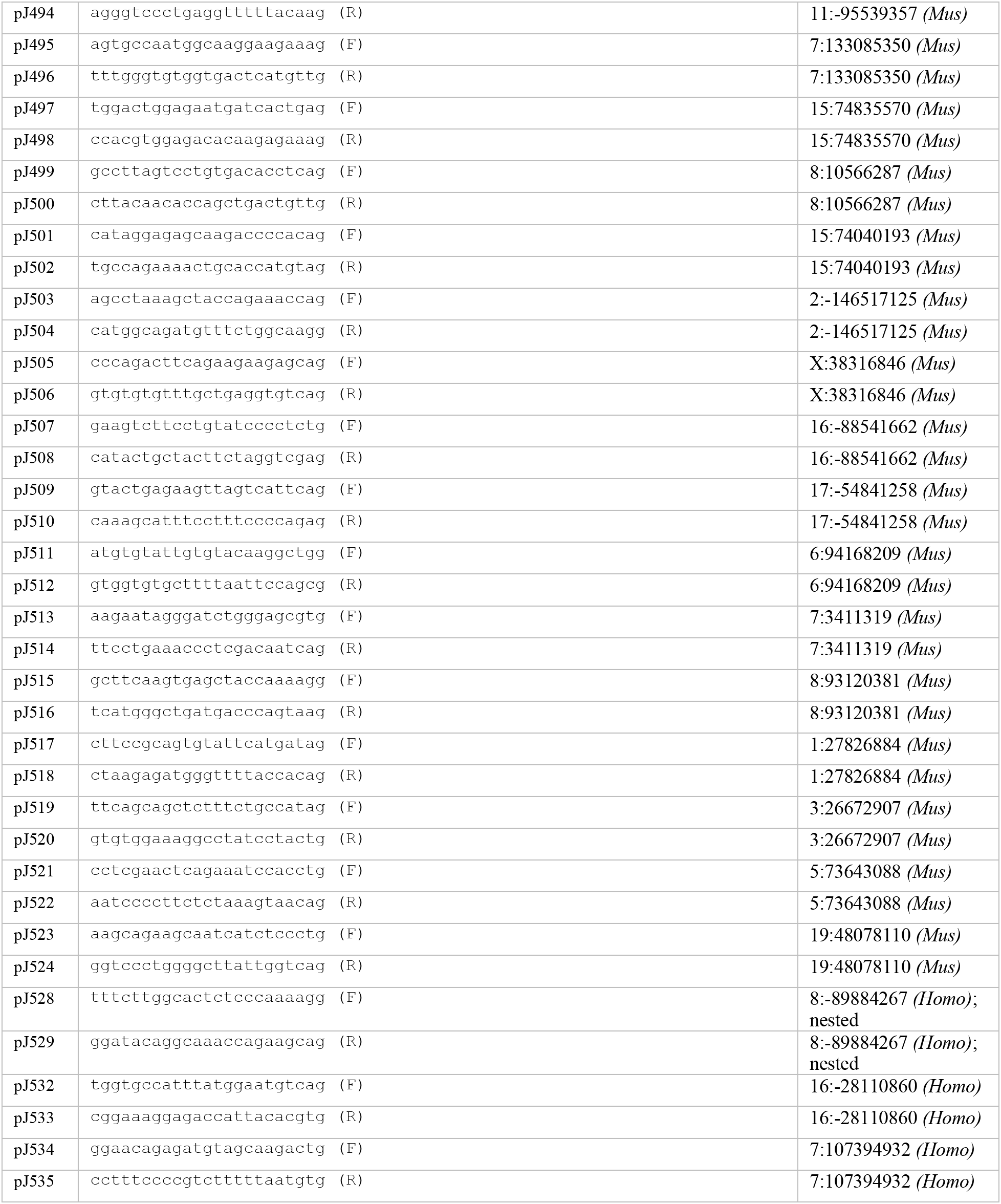
A list of primers and oligonucleotides used in the study.

## REFERENCES

Adegbola, A., Musante, L., Callewaert, B., Maciel, P., Hu, H., Isidor, B., Picker-Minh, S., Le Caignec, C., Delle Chiaie, B., Vanakker, O., et al. 2015). Redefining the MED13L syndrome. Eur J Hum Genet. 23, 1308–1317. 10.1038/ejhg.2015.26.

Aida, T., Nakade, S., Sakuma, T., Izu, Y., Oishi, A., Mochida, K., Ishikubo, H., Usami, T., Aizawa, H., Yamamoto, T., & Tanaka, K. (2016). Gene cassette knock-in in mammalian cells and zygotes by enhanced MMEJ. BMC genomics, 17(1), 979. 10.1186/s12864-016-33.

Anzalone, A. V., Randolph, P. B., Davis, J. R., Sousa, A. A., Koblan, L. W., Levy, J. M., Chen, P. J., Wilson, C., Newby, G. A., Raguram, A., & Liu, D. R. (2019). Search-and-replace genome editing without double-strand breaks or donor DNA. Nature 576, 149–157. 10.1038/s41586-019-1711-4.

Asadollahi, R., Oneda, B., Sheth, F., Azzarello-Burri, S., Baldinger, R., Joset, P., Latal, B., Knirsch, W., Desai, S., Baumer, A., Houge, G., Andrieux, J., & Rauch, A. (2013). Dosage changes of MED13L further delineate its role in congenital heart defects and intellectual disability. Eur J Hum Genet. 21, 1100–1104. 10.1038/ejhg.2013.17.

Asadollahi, R., Zweier, M., Gogoll, L., Schiffmann, R., Sticht, H., Steindl, K., & Rauch, A. (2017). Genotype-phenotype evaluation of MED13L defects in the light of a novel truncating and a recurrent missense mutation. Eur J Med Genet. 60, 451–464. 10.1016/j.ejmg.2017.06.004.

Ayabe, S., Nakashima, K., & Yoshiki, A. (2019). Off- and on-target effects of genome editing in mouse embryos. J Reprod Dev. 65,1–5. 10.1262/jrd.2018-128.

Becher, B., Waisman, A., & Lu, L. F. (2018). Conditional Gene-Targeting in Mice: Problems and Solutions. Immunity, 48, 835–836. 10.1016/j.immuni.2018.05.002.

Cafiero, C., Marangi, G., Orteschi, D., Ali, M., Asaro, A., Ponzi, E., Moncada, A., Ricciardi, S., Murdolo, M., Mancano, G., Contaldo, I., Leuzzi, V., Battaglia, D., Mercuri, E., Slavotinek, A. M., & Zollino, M. (2015). Novel de novo heterozygous loss-of-function variants in MED13L and further delineation of the MED13L haploinsufficiency syndrome. Eur J Hum Genet. 23, 1499–1504. 10.1038/ejhg.2015.19.

Ceccaldi, R., Rondinelli, B., & D’Andrea, A. D. (2016). Repair Pathway Choices and Consequences at the Double-Strand Break. Trends Cell Biol 26, 52–64. 10.1016/j.tcb.2015.07.009.

Chang, H., Pannunzio, N. R., Adachi, N., & Lieber, M. R. (2017). Non-homologous DNA end joining and alternative pathways to double-strand break repair. Nat Rev Mol Cell Biol. 18, 495–506. 10.1038/nrm.2017.48.

Chang, K. T., Jezek, J., Campbell, A. N., Stieg, D. C., Kiss, Z. A., Kemper, K., Jiang, P., Lee, H. O., Kruger, W. D., van Hasselt, P. M., & Strich, R. (2022). Aberrant cyclin C nuclear release induces mitochondrial fragmentation and dysfunction in *MED13L* syndrome fibroblasts. iScience. 25, 103823. 10.1016/j.isci.2022.103823

Chen, H., Choi, J., & Bailey, S. (2014). Cut site selection by the two nuclease domains of the Cas9 RNA-guided endonuclease. J Biol Chem. 289, 13284–13294. 10.1074/jbc.M113.539726.

Chen, J. M., Cooper, D. N., Chuzhanova, N., Férec, C., & Patrinos, G. P. (2007). Gene conversion: mechanisms, evolution and human disease. Nat Rev Genet. 8, 762–775. 10.1038/nrg2193.

Chen, S., Sun, S., Moonen, D., Lee, C., Lee, A. Y., Schaffer, D. V., & He, L. (2019). CRISPR-READI: Efficient Generation of Knockin Mice by CRISPR RNP Electroporation and AAV Donor Infection. Cell Rep. 27(13), 3780–3789.e4. 10.1016/j.celrep.2019.05.103.

Cohen, S. N., Chang, A. C., Boyer, H. W., & Helling, R. B. (1973). Construction of biologically functional bacterial plasmids in vitro. Proc Natl Acad Sci U S A 70, 3240–3244. 10.1073/pnas.70.11.3240.

Conant D, Hsiau T, Rossi N, Oki J, Maures T, Waite K, Yang J, Joshi S, Kelso R, Holden K, Enzmann BL, Stoner R. (2022). Inference of CRISPR Edits from Sanger Trace Data. CRISPR J. 5(1), 123–130. 10.1089/crispr.2021.0113.

Cong, L., Ran, F. A., Cox, D., Lin, S., Barretto, R., Habib, N., Hsu, P. D., Wu, X., Jiang, W., Marraffini, L. A., & Zhang, F. (2013). Multiplex genome engineering using CRISPR/Cas systems. Science 339, 819–823. 10.1126/science.1231143.

Doench, J. G., Hartenian, E., Graham, D. B., Tothova, Z., Hegde, M., Smith, I., Sullender, M., Ebert, B. L., Xavier, R. J., & Root, D. E. (2014). Rational design of highly active sgRNAs for CRISPR-Cas9-mediated gene inactivation. Nat Biotechnol. 32, 1262–1267. 10.1038/nbt.3026.

Doudna J. A. (2020). The promise and challenge of therapeutic genome editing. Nature 578, 229–236. 10.1038/s41586-020-1978-5.

Gaudelli, N. M., Komor, A. C., Rees, H. A., Packer, M. S., Badran, A. H., Bryson, D. I., & Liu, D. R. (2017). Programmable base editing of A•T to G•C in genomic DNA without DNA cleavage. Nature 551, 464–471. 10.1038/nature24644.

Harpak, A., Lan, X., Gao, Z., & Pritchard, J. K. (2017). Frequent nonallelic gene conversion on the human lineage and its effect on the divergence of gene duplicates. Proc Natl Acad Sci U S A 114,12779–12784. 10.1073/pnas.1708151114.

Hay, E. A., Khalaf, A. R., Marini, P., Brown, A., Heath, K., Sheppard, D., & MacKenzie, A. (2017). An analysis of possible off target effects following CAS9/CRISPR targeted deletions of neuropeptide gene enhancers from the mouse genome. Neuropeptides 64, 101–107. 10.1016/j.npep.2016.11.003.

Hsu, P. D., Scott, D. A., Weinstein, J. A., Ran, F. A., Konermann, S., Agarwala, V., Li, Y., Fine, E. J., Wu, X., Shalem, O., Cradick, T. J., Marraffini, L. A., Bao, G., & Zhang, F. (2013). DNA targeting specificity of RNA-guided Cas9 nucleases. Nat Biotechnol. 31, 827–832. 10.1038/nbt.2647.

Iyer, S., Suresh, S., Guo, D., Daman, K., Chen, J., Liu, P., Zieger, M., Luk, K., Roscoe, B. P., Mueller, C., King, O. D., Emerson, C. P., Jr, & Wolfe, S. A. (2019). Precise therapeutic gene correction by a simple nuclease-induced double-stranded break. Nature 568, 561–565. 10.1038/s41586-019-1076-8.

Jiang, P., Song, J., Gu, G., Slonimsky, E., Li, E., & Rosenthal, N. (2002). Targeted deletion of the MLC1f/3f downstream enhancer results in precocious MLC expression and mesoderm ablation. Dev Biol. 243(2), 281–293. doi:10.1006/dbio.2002.0574.

Jinek, M., Chylinski, K., Fonfara, I., Hauer, M., Doudna, J. A., & Charpentier, E. (2012). A programmable dual-RNA-guided DNA endonuclease in adaptive bacterial immunity. Science 337, 816–821. 10.1126/science.1225829.

Kim, D., Kim, J., Hur, J. K., Been, K. W., Yoon, S. H., & Kim, J. S. (2016). Genome-wide analysis reveals specificities of Cpf1 endonucleases in human cells. Nat Biotechnol. 34, 863–868. /10.1038/nbt.3609.

Kim, Y. B., Komor, A. C., Levy, J. M., Packer, M. S., Zhao, K. T., & Liu, D. R. (2017). Increasing the genome-targeting scope and precision of base editing with engineered Cas9-cytidine deaminase fusions. Nat Biotechnol. 35, 371–376. 10.1038/nbt.3803.

Kleinstiver, B. P., Tsai, S. Q., Prew, M. S., Nguyen, N. T., Welch, M. M., Lopez, J. M., McCaw, Z. R., Aryee, M. J., & Joung, J. K. (2016). Genome-wide specificities of CRISPR-Cas Cpf1 nucleases in human cells. Nat Biotechnol. 34, 869–874. 10.1038/nbt.3620.

Klompe, S. E., Vo, P., Halpin-Healy, T. S., & Sternberg, S. H. (2019). Transposon-encoded CRISPR-Cas systems direct RNA-guided DNA integration. Nature 571, 219–225. 10.1038/s41586-019-1323-z.

Li, H., Beckman, K.A., Pessino, V., Huang, B., Weissman, J.S., and Leonetti, M.D. (2017). Design and specificity of long ssDNA donors for CRISPR-based knock-in. bioRxiv. https://doi.org/10.1101/178905.

Lieber, M. R. (2010). The mechanism of double-strand DNA break repair by the nonhomologous DNA end-joining pathway. Annu Rev Biochem. 79,181–211. 10.1146/annurev.biochem.052308.093131.

Mali, P., Aach, J., Stranges, P. B., Esvelt, K. M., Moosburner, M., Kosuri, S., Yang, L., & Church, G. M. (2013). CAS9 transcriptional activators for target specificity screening and paired nickases for cooperative genome engineering. Nat Biotechnol. 31,833–838. 10.1038/nbt.2675.

Meselson, M., & Yuan, R. (1968). DNA restriction enzyme from E. coli. Nature 217, 1110–1114. 10.1038/2171110a0.

Miura, H., Quadros, R. M., Gurumurthy, C. B., & Ohtsuka, M. (2018). Easi-CRISPR for creating knock-in and conditional knockout mouse models using long ssDNA donors. Nat Protoc. 13, 195–215. 10.1038/nprot.2017.153.

Paquet D, Kwart D, Chen A, Sproul A, Jacob S, Teo S, Olsen KM, Gregg A, Noggle S, Tessier-Lavigne M. (2016). Efficient introduction of specific homozygous and heterozygous mutations using CRISPR/Cas9. Nature. 533, 125–129. 10.1038/nature17664

Pawelczak, K. S., Gavande, N. S., VanderVere-Carozza, P. S., & Turchi, J. J. (2018). Modulating DNA Repair Pathways to Improve Precision Genome Engineering. ACS Chem Biol. 13, 389–396. 10.1021/acschembio.7b00777.

Ran, F. A., Cong, L., Yan, W. X., Scott, D. A., Gootenberg, J. S., Kriz, A. J., Zetsche, B., Shalem, O., Wu, X., Makarova, K. S., Koonin, E. V., Sharp, P. A., & Zhang, F. (2015). In vivo genome editing using Staphylococcus aureus Cas9. Nature 520, 186–191. 10.1038/nature14299.

Ran, F. A., Hsu, P. D., Lin, C. Y., Gootenberg, J. S., Konermann, S., Trevino, A. E., Scott, D. A., Inoue, A., Matoba, S., Zhang, Y., & Zhang, F. (2013). Double nicking by RNA-guided CRISPR Cas9 for enhanced genome editing specificity. Cell 154, 1380–1389. 10.1016/j.cell.2013.08.021.

Smol, T., Petit, F., Piton, A., Keren, B., Sanlaville, D., Afenjar, A., Baker, S., Bedoukian, E. C., Bhoj, E. J., Bonneau, D., et al. (2018). MED13L-related intellectual disability: involvement of missense variants and delineation of the phenotype. Neurogenetics 19, 93–103. 10.1007/s10048-018-0541-0.

Strecker, J., Ladha, A., Gardner, Z., Schmid-Burgk, J. L., Makarova, K. S., Koonin, E. V., & Zhang, F. (2019). RNA-guided DNA insertion with CRISPR-associated transposases. Science 365, 48–53. 10.1126/science.aax9181.

Thomas, K. R., & Capecchi, M. R. (1987). Site-directed mutagenesis by gene targeting in mouse embryo-derived stem cells. Cell 51, 503–512. 10.1016/0092-8674(87)90646-5.

Truong, L. N., Li, Y., Shi, L. Z., Hwang, P. Y., He, J., Wang, H., Razavian, N., Berns, M. W., & Wu, X. (2013). Microhomology-mediated End Joining and Homologous Recombination share the initial end resection step to repair DNA double-strand breaks in mammalian cells. Proc Natl Acad Sci U S A 110, 7720–7725. 10.1073/pnas.1213431110.

van Haelst, M. M., Monroe, G. R., Duran, K., van Binsbergen, E., Breur, J. M., Giltay, J. C., & van Haaften, G. (2015). Further confirmation of the MED13L haploinsufficiency syndrome. Eur J Hum Genet. 23, 135–138. 10.1038/ejhg.2014.69.

Wang, H., Yang, H., Shivalila, C. S., Dawlaty, M. M., Cheng, A. W., Zhang, F., & Jaenisch, R. (2013). One-step generation of mice carrying mutations in multiple genes by CRISPR/Cas-mediated genome engineering. Cell 153, 910–918. 10.1016/j.cell.2013.04.025.

Yamamoto, Y., & Gerbi, S. A. (2018). Making ends meet: targeted integration of DNA fragments by genome editing. Chromosoma 127, 405–420. 10.1007/s00412-018-0677-6.

Yang, H., Wang, H., Shivalila, C. S., Cheng, A. W., Shi, L., & Jaenisch, R. (2013). One-step generation of mice carrying reporter and conditional alleles by CRISPR/Cas-mediated genome engineering. Cell 154, 1370–1379. 10.1016/j.cell.2013.08.022.

Yokouchi, Y., Suzuki, S., Ohtsuki, N., Yamamoto, K., Noguchi, S., Soejima, Y., Goto, M., Ishioka, K., Nakamura, I., Suzuki, S., Takenoshita, S., & Era, T. (2020). Rapid repair of human disease-specific single-nucleotide variants by One-SHOT genome editing. Sci Rep. 10, 13927. 10.1038/s41598-020-70401-7.

Yoshimi, K., Kunihiro, Y., Kaneko, T., Nagahora, H., Voigt, B., & Mashimo, T. (2016). ssODN-mediated knock-in with CRISPR-Cas for large genomic regions in zygotes. Nat Commun. 7, 10431. 10.1038/ncomms10431.

Zetsche, B., Gootenberg, J. S., Abudayyeh, O. O., Slaymaker, I. M., Makarova, K. S., Essletzbichler, P., Volz, S. E., Joung, J., van der Oost, J., Regev, A., Koonin, E. V., & Zhang, F. (2015). Cpf1 is a single RNA-guided endonuclease of a class 2 CRISPR-Cas system. Cell 163, 759–771. 10.1016/j.cell.2015.09.038.

